# Host lipids regulate multicellular behavior of a predator of a human pathogen

**DOI:** 10.1101/2024.01.31.578218

**Authors:** Ria Q. Kidner, Eleanor B. Goldstone, Martina R. Laidemitt, Melissa C. Sanchez, Catherine Gerdt, Lorin P. Brokaw, Núria Ros-Rocher, Jamie Morris, W. Sean Davidson, Joseph P. Gerdt

**Affiliations:** Department of Chemistry, Indiana University, Bloomington, IN 47405, USA; Department of Biology, Center for Evolutionary and Theoretical Immunology, Parasite Division, Museum of Southwestern Biology, University of New Mexico, Albuquerque, New Mexico 87131, USA; Department of Functional Genomics and Evolution, Institut de Biologia Evolutiva (Consejo Superior de Investigaciones Científicas-Universitat Pompeu Fabra), Passeig Marítim de la Barceloneta 37-49, 08003 Barcelona, Spain; Department of Cell Biology and Infection and Department of Developmental and Stem Cell Biology, Institut Pasteur, Université Paris-Cité, CNRS UMR3691, 25-28 Rue du Docteur Roux, 75015, Paris, France; Department of Pathology and Laboratory Medicine, University of Cincinnati, Cincinnati OH 45237, USA

## Abstract

As symbionts of animals, microbial eukaryotes benefit and harm their hosts in myriad ways. A model microeukaryote (*Capsaspora owczarzaki*) is a symbiont of *Biomphalaria glabrata* snails and may prevent transmission of parasitic schistosomes from snails to humans. However, it is unclear which host factors determine *Capsaspora’*s ability to colonize snails. Here, we discovered that *Capsaspora* forms multicellular aggregates when exposed to snail hemolymph. We identified a molecular cue for aggregation: a hemolymph-derived phosphatidylcholine, which becomes elevated in schistosome-infected snails. Therefore, *Capsaspora* aggregation may be a response to the physiological state of its host, and it may determine its ability to colonize snails and exclude parasitic schistosomes. Furthermore, *Capsaspora* is an evolutionary model organism whose aggregation may be ancestral to animals. This discovery, that a prevalent lipid induces *Capsaspora* multicellularity, suggests that this aggregation phenotype may be ancient. Additionally, the specific lipid will be a useful tool for further aggregation studies.

## INTRODUCTION

Microbial symbionts frequently impact the fitness of their animal hosts—both for the better and the worse.^1,2^ Due to their abundance and ease of study, bacterial symbionts have garnered the most research. However, microbial *eukaryotes* (*i.e.,* protists) also frequently influence their hosts.^3^ Although pathogens are the best studied eukaryotic symbionts (*e.g., Plasmodium*, *Leishmania*, *Candida*, and chytrids),^4–7^ mutualist and commensal microeukaryotes also populate the literature.^8–10^ The protist *Capsaspora owczarzaki* (hereafter “*Capsaspora*”) is an intriguing symbiont of snails that may *both* reveal insight into protist-animal symbioses *and* curtail the spread of neglected tropical diseases.^11,12^

*Capsaspora* was initially isolated as unicellular filopodiated amoebae from the pericardia and mantles of *Biomphalaria glabrata*. This snail is also the intermediate host that transmits *Schistosoma mansoni,* the causative agent of intestinal human schistosomiasis in Africa and the Neotropics (**Fig. 1A**).^11,12^ Due to its disease relevance, *Biomphalaria* snails have been well studied in the laboratory for decades.^13–17^ More recently, *Capsaspora* has also become an emerging experimental model with substantial “omic” resources^18–23^ and molecular tools available,^24–27^ making this snail-amoeba symbiosis ideally suited for deeper analysis as a model system. More significantly, *Capsaspora* can readily adhere to and kill schistosomes while they are sporocysts (the intramolluscan growth stage) *in vitro*.^12,28^ Therefore, *Capsaspora* may be able to halt the spread of schistosomiasis by outcompeting schistosomes within their intermediate host snails— similar to how *Wolbachia* bacteria halt the spread of mosquito-transmitted diseases.^29–31^ Since the ∼300 million people who suffer snail-transmitted diseases can be difficult to treat, such an ecological intervention to deplete parasites in endemic areas is an attractive approach.^32–34^ However, the interactions between *Capsaspora*, *Biomphalaria* snails, and schistosome parasites are still poorly understood.^11,12,28,35^

**Figure 1.**
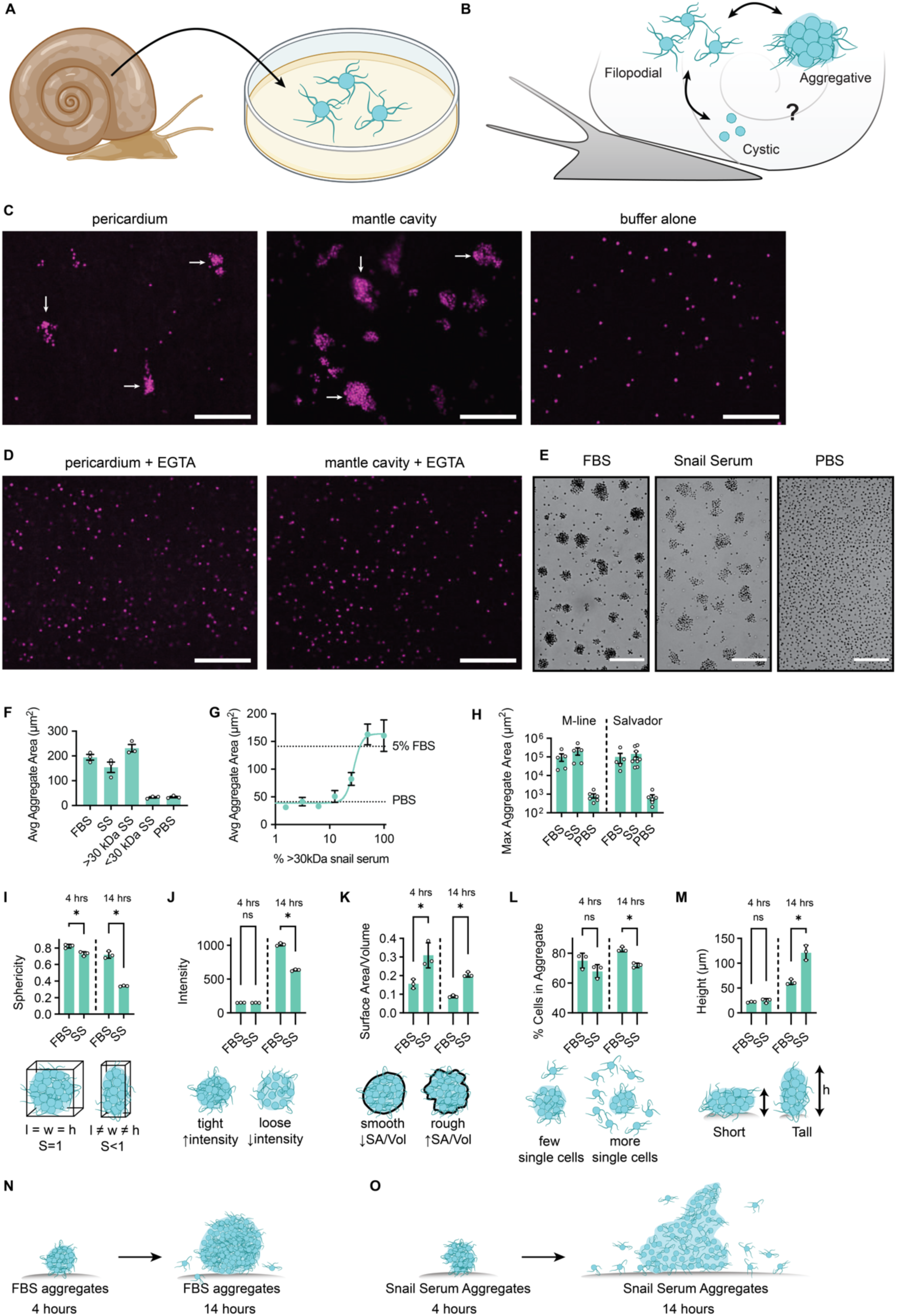
*Capsaspora* aggregates in response to snail tissue and serum. (A) *Capsaspora* was originally isolated from the pericardium and mantle of *B. glabrata*. Cells that grew out from snail samples were filopodiated. (B) Known *Capsaspora* life stages: filopodiated amoebae, cysts, and multicellular aggregates. It was previously unclear which life stages of *Capsaspora* are present inside snails. (C) Representative images of tdTomato-expressing *Capsaspora* (ATCC®30864) aggregates observed after injection into *B. glabrata* (NMRI) snail tissues (left and center images) compared to a negative control where *Capsaspora* was injected directly onto a microscope slide with no snail (right image). White arrows indicate example aggregates. (D) Representative images of *Capsaspora* co-injected with a calcium chelator (EGTA, 250 mM) into snail tissues—aggregation was prevented. (E) Representative images of *Capsaspora* aggregates induced by either 5% (v/v) FBS or 50% (v/v) snail serum compared to cells treated with 5% (v/v) 1X PBS buffer negative control. (F) Average area of cell aggregates induced by FBS, snail serum (SS), and small molecules (<30 kDa) and macromolecules (>30 kDa) from snail serum. (G) Dilution series of >30 kDa snail serum shows a dose-dependent induction of aggregation with an EC50 ∼30% (v/v). (H) Area of aggregates induced by either 10% (v/v) FBS or 50% (v/v) >30 kDa snail serum in multiple isolated strains of *Capsaspora*. All strains tested showed aggregation in response to snail serum. (I–M) Analysis of 3D confocal microscopy images of *Capsaspora* cells after 4 hours and 14 hours of induction with either 5% (v/v) FBS or 50% (v/v) snail serum. Cells at 4 hours were tdTomato-expressing *Capsaspora*, and cells at 14 hours were stained with 0.02 mg/mL propidium iodide. (I) Sphericity of aggregates, determined by the ratio of aggregate bounding box dimensions. The average snail serum-induced aggregate was less spherical than FBS-induced aggregates. (J) The density of snail serum aggregates, calculated by the average intensity of stained-cell fluorescence, was similar to the density of FBS aggregates at 4 hours and lower than FBS aggregates at 14 hours. (K) The roughness of the aggregate surface, calculated by the surface area to volume ratio, was higher in snail serum aggregates at both time points. (L) The percentage of cells included within an aggregate, relative to the total number of cells in an image, showed there were slightly more non-aggregated cells present in snail serum-induced samples compared to those with FBS induction. This effect was more significant at 14 hours than 4 hours. (M) The average heights of the snail serum-induced aggregates were similar to the average FBS-induced aggregates at 4 hours but reached substantially taller by 14 hours. (N) Cartoons representing side views of typical aggregates induced by 5% (v/v) FBS at 4 hours and at 14 hours. (O) Cartoons representing side views of typical aggregates induced by 50% (v/v) snail serum at 4 hours and at 14 hours. Representative images of FBS-induced aggregates and snail serum-induced aggregates from a top-view and a side-view at 4 hours and 14 hours are shown in **Fig. S2A–D**. For microscopy images, scale bars are 100 µm. For plots, mean ± sem (n=3) are shown, and values from individual replicates are displayed with small circles.

Although *Capsaspora* has been isolated from multiple inbred lines of *B. glabrata*^11^ and molecularly detected by sequencing from wild snails,^36^ it remains absent from many *B. glabrata* snails in the laboratory and the wild. Moreover, it is unknown which host factors determine *Capsaspora*’s ability to colonize the snail, and it is also unclear what fitness impact *Capsaspora* has on its host snail and co-resident parasites. Additionally, multiple life stages of *Capsaspora* have been described in the laboratory,^23^ but it is unclear which life stages are relevant to its behavior inside the host snail (**Fig. 1B**). In this study, we aimed to determine if *Capsaspora* could respond to chemical or cellular factors in its host snail environment.

We found that within snail host tissue, *Capsaspora* formed multicellular aggregates. These aggregates appeared similar to those formed by *Capsaspora* upon *in vitro* exposure to fetal bovine serum.^27^ Furthermore, we discovered that the aggregation inducer in the snail serum is a specific phosphatidylcholine lipid (or possibly a suite of phosphatidylcholine lipids). Remarkably, the concentration of this lipid in the snail hemolymph increased when the snail was infected with schistosomes, which led to significantly greater *Capsaspora* aggregation. *Capsaspora* also aggregated differently in hemolymph from different inbred snail lines. Therefore, *Capsaspora* can sense and respond to the physiological state and identity of its host snail. This work raises the hypothesis that a chemical mechanism of host discrimination may explain the presence of *Capsaspora* in some snails but not others. Further dissection of its *in vivo* aggregation and persistence may reveal the molecular requirements for *Capsaspora* to colonize its host and possibly exclude pathogenic schistosomes.

## RESULTS

### *Capsaspora* forms multicellular aggregates in snail tissue

To obtain an initial glimpse into the interaction of *Capsaspora* with its host snail, we introduced fluorescently labeled *Capsaspora* cells in the filopodial stage into naïve *Biomphalaria glabrata* (NMRI) snails. The snails had no prior schistosome infection or colonization with *Capsaspora* (as evidenced by PCR, see *Supporting Information* **Fig. S1A–B**). Since *Capsaspora* has been isolated from snail pericardial explants and mantle explants and swabs,^12^ we injected *Capsaspora* cells into the pericardia and mantles. We discovered that *Capsaspora* formed multicellular aggregates within 5 minutes of injection into the tissue (**Fig. 1C**, left and center images, **Fig. S1C– D**). In contrast, *Capsaspora* imaged in the absence of snail tissue (in the injection buffer control) failed to aggregate (**Fig. 1C**, right image), verifying that components of the snail triggered *Capsaspora* aggregation. Notably, these aggregates were reminiscent of the aggregates observed upon addition of fetal bovine serum (FBS) to *Capsaspora* cells *in vitro.*^27^ Like FBS-induced aggregation, this aggregation phenotype was calcium dependent: co-injection of excess EGTA (a calcium-specific chelator) with *Capsaspora* into the snail pericardia and mantles suppressed the aggregation phenotype (**Fig. 1D**). The observation of *Capsaspora* aggregation inside its natural snail host suggests that *Capsaspora*’s previously-observed aggregation phenotype may be ecologically relevant within the natural snail environment (not an artefact of artificial *in vitro* growth media).

### *Capsaspora* forms multicellular aggregates in response to macromolecules in host snail serum

*Capsaspora* was also previously isolated from snail hemolymph.^11^ Therefore, we asked whether hemolymph alone could induce *Capsaspora* aggregation *in vitro*. Indeed, sterile-filtered (0.22 µm) hemolymph from NMRI snails (hereafter “snail serum” or “SS”) induced cellular aggregation (**Fig. 1E–F**). Because the aggregate inducer previously found in FBS was a large lipoprotein complex,^27^ we hypothesized that the inducer in snail serum was a macromolecule. To test this hypothesis, we fractionated snail serum using a 30 kDa MW cutoff filter, and the large and small fractions were tested separately. As hypothesized, we found that >30 kDa snail serum induced robust aggregation, and the <30 kDa fraction did not (**Fig. 1F**). Aggregation induced by >30 kDa snail serum was dose-dependent with an EC50 ∼30% (v/v) (**Fig. 1G**).

We also asked if the behavior was conserved across multiple isolates of *Capsaspora* (or if it was possibly a phenotype unique to a single *Capsaspora* strain). In addition to the well-studied ATCC 30864 *Capsaspora* strain, we tested the ability of snail serum to induce aggregation of two other *Capsaspora* strains isolated separately from M-line and Salvador *B. glabrata* snails (ATCC 50973 and 50974, respectively).^11^ Both of the alternative *Capsaspora* strains aggregated in response to snail serum, as well as FBS (**Fig. 1H**). Notably, the other two cultures are less studied and grow slower *in vitro*, possibly indicating less adaptation to laboratory culture conditions. Therefore, we believe that the aggregative response of *Capsaspora* to *Biomphalaria* serum components is likely a widespread and natural *Capsaspora* phenotype. The original ATCC 30864 strain was used for further studies in this manuscript.

Compared to previous reports of *Capsaspora* aggregation, we noticed that the snail serum-induced aggregates appeared less circular and less dense than those induced by FBS. This observation motivated us to characterize the aggregation morphology of *Capsaspora* cells induced by snail serum compared to FBS-induced aggregates via confocal microscopy. We assessed both early and mature aggregates (4 hours and 14 hours after addition of serum). Indeed, we found that snail serum-induced aggregates were quantitatively less spherical than those induced by FBS, as measured by the ratio of bounding box dimensions of each aggregate (**Fig. 1I**). The difference was most noticeable in mature 14-hour aggregates. Regarding cell density (measured by the intensity of cell fluorescence within the boundary of each aggregate), snail serum-induced aggregates were initially similar to FBS-induced aggregates at 4 hours but were less dense by 14 hours (**Fig. 1J**). Additionally, snail serum-induced aggregates were rougher around the edges than those induced by FBS, causing a higher surface area to volume ratio at both time points (**Fig. 1K**). Furthermore, snail serum-induced samples had more individual cells not encompassed in aggregates than the FBS-induced samples. This difference was significant at the 14-hour time point (**Fig. 1L**). Finally, while the snail serum-induced aggregates were of similar height to FBS-induced aggregates at 4 hours, they ultimately reached greater height than those induced by FBS at 14 hours (yet, this measurement was biased by especially tall spires in the more asymmetric snail serum aggregates) (**Fig. 1M–O and Fig. S2A–D)**. Interestingly, the morphological differences observed here under different chemical stimulation mirror some of the morphological differences reported in mutant *Capsaspora* strains.^26,37^ Therefore, it is likely that multiple chemical and genetic factors contribute to the specific multicellular structures formed by *Capsaspora* cell-cell adhesion.

In parallel, we also monitored the kinetics of cellular aggregation in response to snail serum and compared it with aggregates induced using FBS. Although FBS-induced aggregates lasted slightly longer than snail serum-induced aggregates, both persisted for over 20 hours. (**Fig. 2A– B** and **Movie S1A–B**). We observed a small spike in aggregate size in the snail serum samples at early time points. This was due to the initial formation of large, less circular aggregates, that later divided into several smaller and rounder aggregates (**Fig. 2B–C**). This spike was not observed in the FBS sample, because the aggregates became circular much more quickly. Overall, snail serum and FBS induced similar—but not identical—aggregation morphology and kinetics.

**Figure 2:**
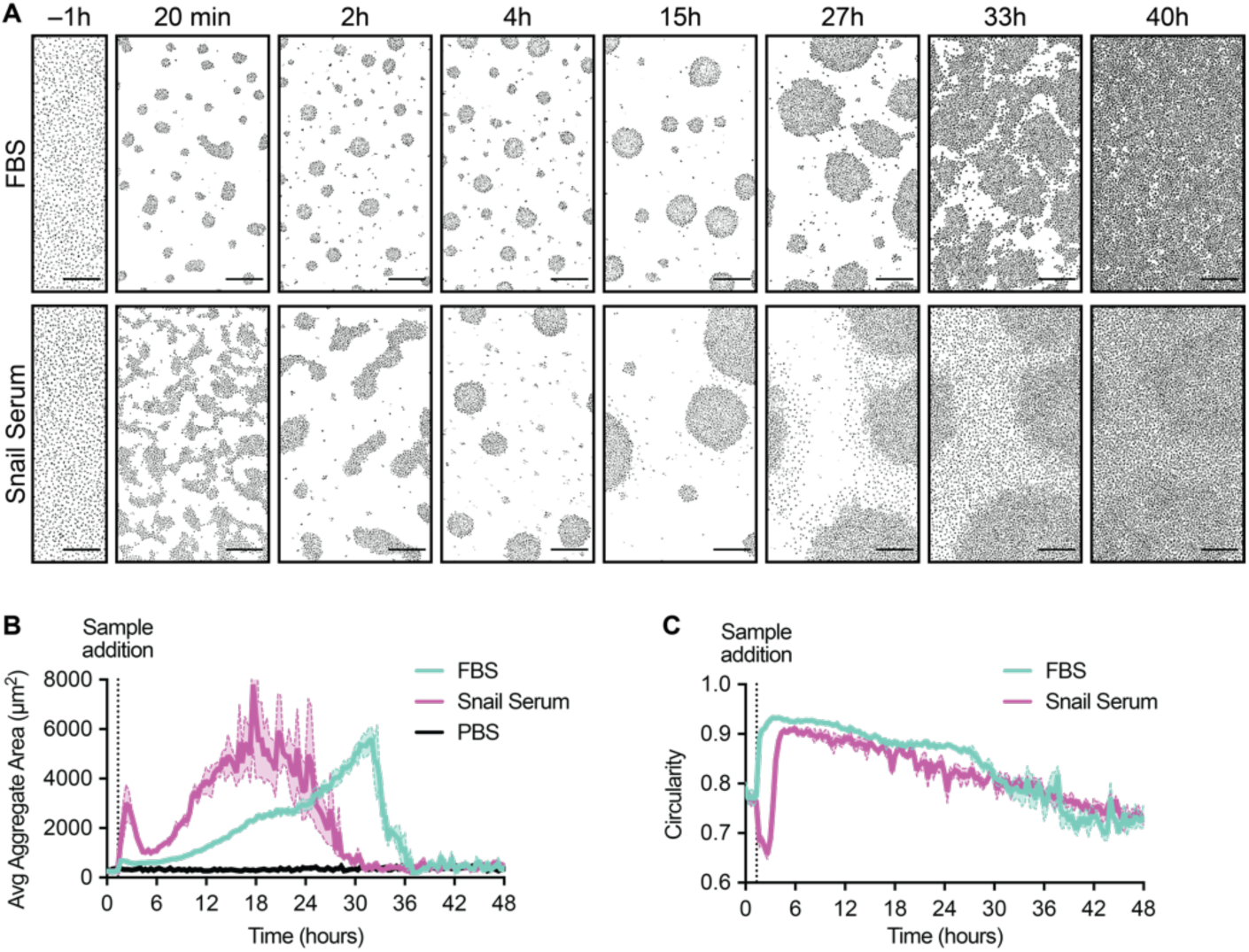
Dynamics of Capsaspora aggregates over time. (A) Representative images showing cellular aggregates monitored for 2 days after addition of 5% (v/v) FBS or 50% (v/v) >30 kDa snail serum components. Images were converted to binary in FIJI to enhance contrast. Scale bar is 250 µm. The full time-lapse is available as **Movie S1**. (B) The average aggregate areas measured every 20 minutes for 48 hours after addition of 5% (v/v) FBS or 50% (v/v) >30 kDa snail serum components. FBS-induced aggregates gradually increased in area over ∼30 hours followed by a sudden disaggregation; however, snail serum induced an initial spike where aggregates were amorphous followed by the formation of circular aggregates that gradually enlarged and then gradually disaggregated. (C) Average circularity of aggregates calculated in two dimensions measured every 20 minutes for 48 hours after addition of 5% (v/v) FBS or 50% (v/v) >30 kDa snail serum components shows initial snail serum-induced aggregates were amorphous. Plots show means ± sem (n=3).

Together, these results suggest that serum-induced cellular aggregation is a relevant response of *Capsaspora* to its host snail environment. We next sought to determine the specific identity of the snail serum inducer(s).

### Lipids extracted from snail serum are sufficient to trigger aggregation

Because protein-free lipid particles were previously shown to induce aggregation,^27^ we hypothesized that the macromolecular aggregation inducer(s) in snail serum were also lipid complexes. To test this hypothesis, we extracted the total lipids from snail serum and tested the solubilized lipid extract for aggregation induction. Indeed, we found that the crude lipids induced robust aggregation activity with an EC50 around 100 µg/mL (**Fig. 3A**). This concentration was similar to the concentration of total lipids present in active dilutions of >30 kDa snail serum (see **Materials and Methods** section), suggesting that lipids are sufficient to account for the aggregation induction activity in >30 kDa snail serum.

**Figure 3.**
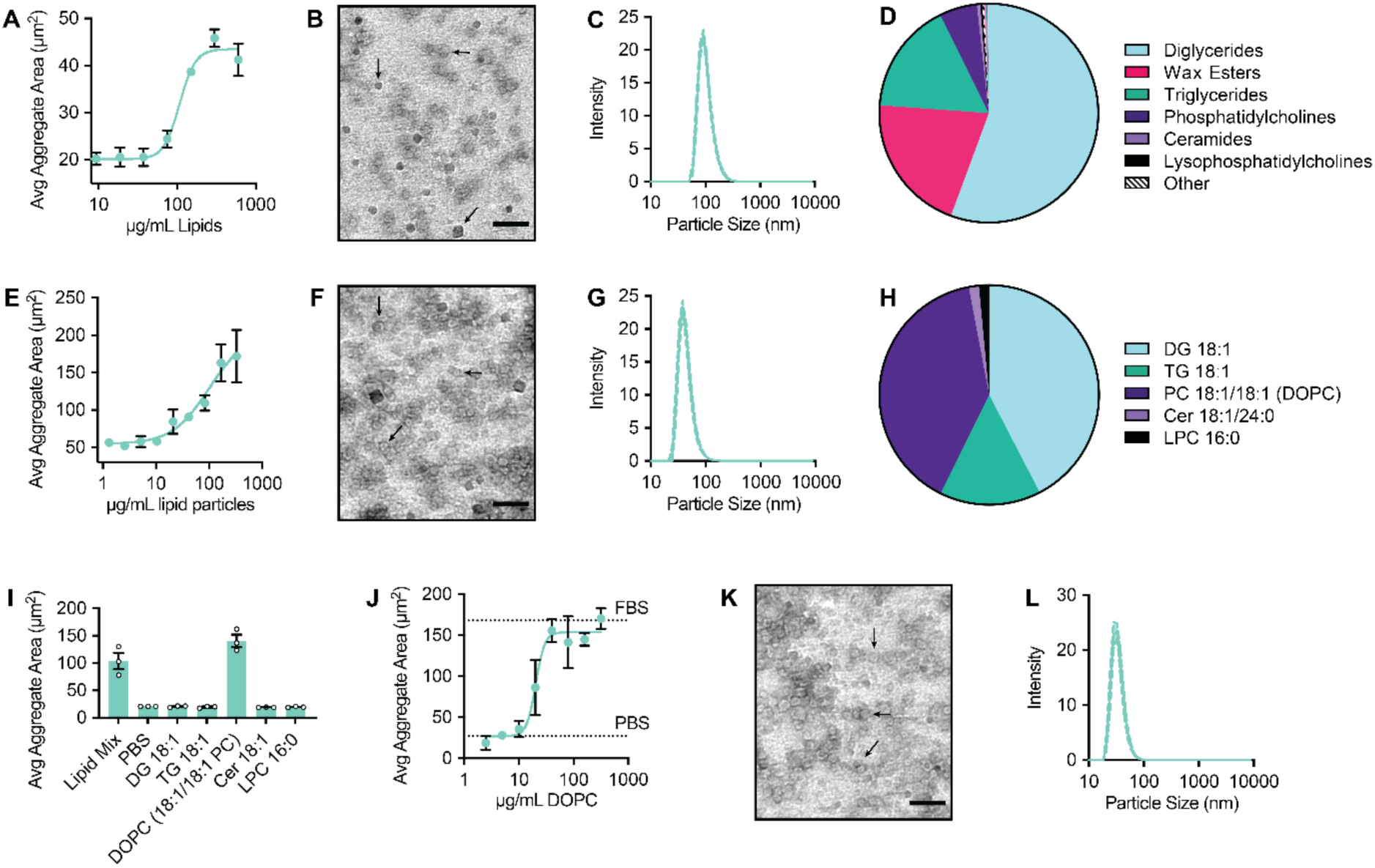
Lipids isolated from *Biomphalaria* snail serum induce aggregation. (A) Average area of aggregates induced by a dilution series of lipids isolated from snail serum and prepared into soluble emulsions. Area plotted as a function of lipid concentration in µg/mL. Lipids induced aggregation in a dose-dependent manner with an EC50 = 100 µg/mL. (B) Representative TEM image of prepared crude snail serum lipid particles show active lipids were incorporated into particles with a range of sizes. (C) Crude snail serum lipid particles were ∼90 nm in diameter as measured by DLS. (D) Pie chart of the major lipid classes present in the prepared and tested crude snail serum lipid particles. The most abundant lipid classes were diglycerides, wax esters, triglycerides, and phosphatidylcholines. (E) Average area of aggregates induced by a simple mixture of synthetic lipids. The simple mix contained glyceryl dioleate (DG), glyceryl trioleate (TG), dioleoyl phosphatidylcholine (PC), 18:1/24:0 ceramide (Cer), and palmitoyl lysophosphatidylcholine (LPC). Lipids significantly induced aggregation at concentrations ∼100 µg/mL, similarly to the extracted lipids. (F) Representative TEM image of particles prepared using the simple synthetic lipid mix showed a range of sizes, much like the natural lipid preparation. (G) Synthetic lipid mixture particles were ∼50 nm in diameter as measured by DLS. (H) Pie chart of the final lipid ratios included in the soluble synthetic lipid particles. (I) Average area of aggregates induced by either the simple lipid mix or the individual lipids tested at the same concentration as present in the simple lipid mix. Of the individual lipids, only DOPC induced aggregation. (J) Average area of aggregates induced by pure DOPC lipid vesicles. DOPC induced aggregation in a dose-dependent manner (EC50 = 20 µg/mL). (K) Representative TEM image of DOPC vesicles. (L) DOPC vesicles were ∼40 nm in diameter as measured by DLS. For plots, means ± sem (n=3) are shown, and values from individual replicates are displayed with small circles. For microscopy images, scale bars are 100 nm, arrows highlight example particles.

We then physically and chemically characterized these emulsified crude serum lipid particles that induced aggregation. First, we used transmission electron microscopy (TEM) to visualize particle sizes (**Fig. 3B**) and dynamic light scattering (DLS) to determine the particle diameters. The crude snail lipid particles were ∼90 nm (**Fig. 3C**). This value is slightly more than three times the diameter of low density lipoproteins (LDLs, which are the aggregation inducers in FBS).^27^ Next, we employed LC-MS to evaluate the chemical composition of the solubilized snail lipid particles. Although analysis of crude snail serum extracts revealed the presence of over 800 lipids, (see supplementary **Table S1 and Fig. S3**), the solubilized lipid particles did not contain the full array of serum lipids (*i.e.*, some serum lipids resisted resuspension and were not delivered to the cells). Nonetheless, a complex mixture of major lipid classes including diglycerides, wax esters, triglycerides, and phospholipids was clearly incorporated into these aggregate-inducing particles (**Fig. 3D** and **Fig. S3**). Therefore one (or many) lipids present in snail serum induce multicellular aggregation of *Capsaspora*.

We then asked if the aggregation activity was due to a minor component or a major lipid present in the extracted snail serum. First, to determine if a major lipid was responsible for the activity, we prepared a simplified lipid mixture from representative lipids of each of the major lipid classes present in the snail serum extract. The lipid mix, prepared from commercially available synthetic lipids, contained glyceryl dioleate (DG), glyceryl trioleate (TG), dioleoyl phosphatidylcholine (PC), 18:1/24:0 ceramide (Cer), and palmitoyl lysophosphatidylcholine (LPC). Despite being a large class represented in the extracted lipids, we did not include wax esters in this initial simple mix because we could not easily obtain short-tailed wax esters. We combined the five aforementioned lipids at approximately their natural ratio in snail serum and solubilized them by sonication into mixed-lipid particles. Remarkably, this simplified mixture induced aggregation in a dose-dependent manner with similar potency to the crude snail lipids (**Fig. 3E**). We also validated the formation of lipid particles by TEM and DLS measurements as before, which revealed particles that were ∼50 nm in diameter (**Fig. 3F–G**). We also determined the final lipid ratio in the particles by LC-MS to be slightly different than the original intended ratio (**Fig. 3H**), yet all lipids were incorporated into the soluble particles. Thus, a simple mixture of the major snail serum lipids is sufficient to induce *Capsaspora* aggregation.

Then, to determine if a single lipid from snail serum was sufficient to induce aggregation, we tested each component of the active lipid mix individually. We added each lipid to *Capsaspora* (at the same concentration as in the simple lipid mix) and found pure dioleoyl PC (DOPC) lipids to be active, while no other lipids elicited an aggregation response (**Fig. 3I**). The DOPC lipids induced robust aggregation in a dose dependent manner with an EC50 of 20 µg/mL (**Fig. 3J**), which is about five times more potent than the extracted natural lipids from snail serum and the simplified lipid mix. We also validated the formation of lipid vesicles of ∼40 nm diameter with TEM and DLS (**Fig. 3K–L**). Moreover, to determine if 20 µg/mL is a biologically relevant concentration, we quantified the amount of DOPC in snail serum using LC-MS. Based on spectral intensity normalized to an internal PC control, we estimated the concentration of DOPC in snail serum to be about 2 µg/mL (∼10X lower than the EC50 of pure DOPC vesicles). Therefore, DOPC is likely not the sole inducer of aggregation present in snail serum (*e.g.* other PCs in snail serum may contribute as well). In summary, pure DOPC, a lipid present in snail serum, is sufficient to induce *Capsaspora* multicellular aggregation; however, it is likely that other lipids also contribute to the phenotype.

Since the morphologies of FBS- and snail serum-induced aggregates had differed, we next assessed the morphology of DOPC-induced aggregates using the same quantifiable features measured in **Fig. 1**. We found that DOPC-induced aggregates generally resembled those induced by FBS and/or snail serum and that their morphology depended some on the concentration of DOPC (**Fig. 4A–K**). At intermediate DOPC concentrations, the DOPC-induced aggregates were similar in sphericity, cell density, and smoothness to snail serum-induced aggregates (**Fig. 4B– D, G–I, L–N**). However, more cells remained outside of aggregates in the DOPC-induced condition than those induced by either FBS or snail serum (**Fig. 4E, J, O**). The DOPC-induced aggregates were similar in height to FBS-induced aggregates (**Fig. 4F, K, P**). Overall, most morphological features of DOPC-induced aggregates were similar to those induced by snail serum or FBS. The dependence of morphology on the identity and concentration of lipids suggested that *Capsaspora* aggregation morphology may differ depending on the exact hemolymph composition of its host.

**Figure 4.**
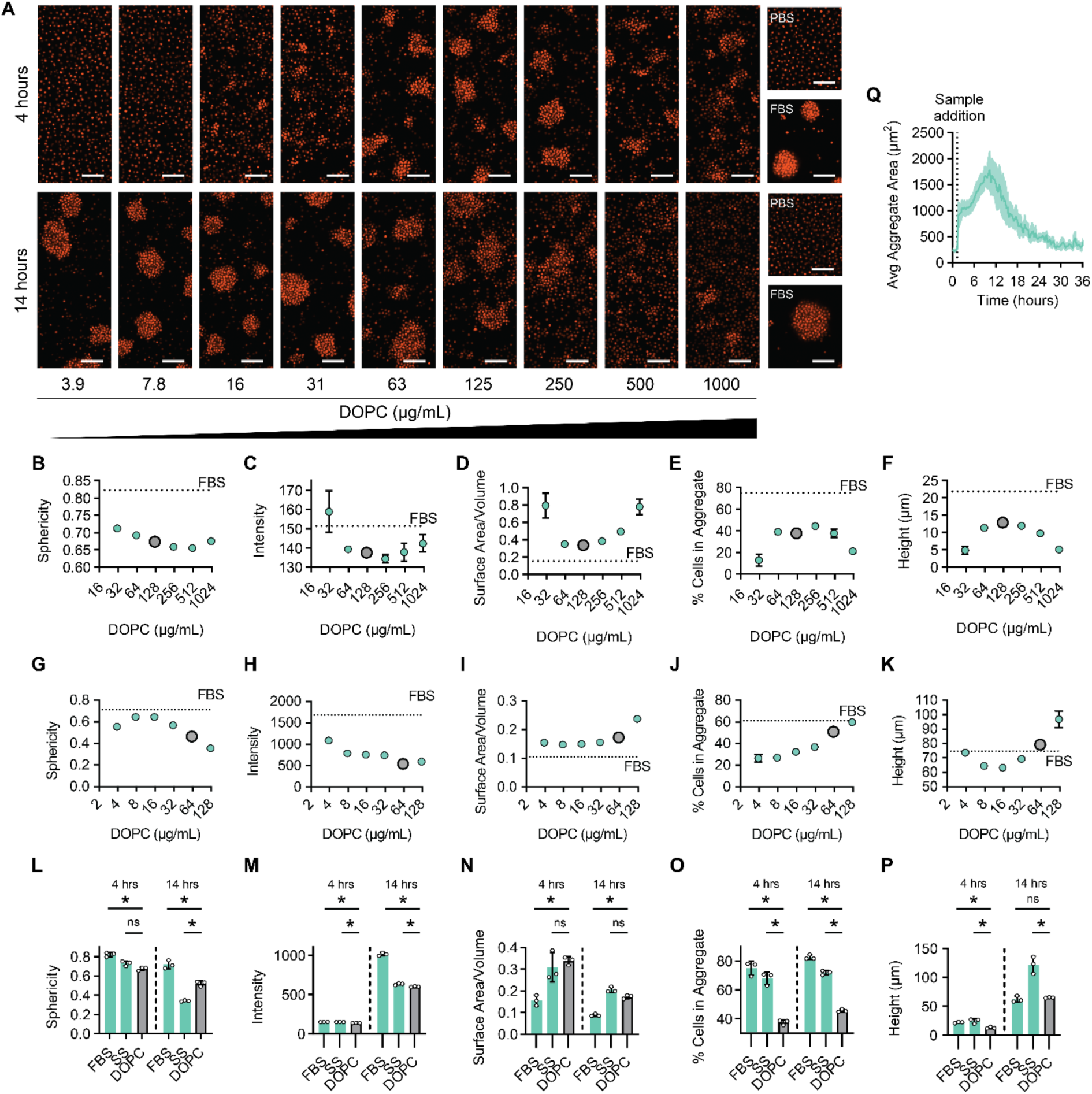
*Capsaspora* aggregate morphology is sensitive to the nature of the inducer, concentration of the inducer, and time. (A) Representative confocal microscopy images of tdTomato-expressing *Capsaspora* cell aggregates after 4 hours and 14 hours of induction with increasing concentrations of DOPC. Scale bars are 50 µm. (B–F) Plots of morphology features of aggregates after 4 hours of induction with increasing concentrations of DOPC. The dotted line represents the value for FBS-induced aggregates, and the larger grey circle is the concentration used in L–P (125 µg/mL for 4 hours). (G–K) Plots of morphology features of aggregates after 14 hours of induction with increasing concentrations of DOPC. The dotted line represents the value for FBS-induced aggregates, and the larger grey circle is the same concentration used in L–P (63 µg/mL for 14 hours). (B/G) Sphericity of aggregates as determined by the ratio of aggregate bounding box dimensions. The sphericity of aggregates generally decreased with increasing concentration of DOPC. (C/H) The density of aggregates induced by DOPC as calculated by the average intensity of fluorescence. The density generally decreased with increasing DOPC concentration. (D/I) The roughness of the aggregate surface calculated by the surface area to volume ratio. The roughness of aggregates increased at the highest concentrations of DOPC. (E/J) The percentage of cells included in an aggregate compared to the total number of cells in the image positively correlated with the concentration of DOPC at 14 hours, but exhibited a non-monotonic dose response at 4 hours. (F/K) The height of aggregates measured by analysis of confocal microscopy images in three dimensions. At 14 hours, height positively correlated with the concentration of DOPC, but again a non-monotonic dose response was observed at 4 hours. (L–P) Plots of morphology features of aggregates after 4 hours and 14 hours of induction with either 5% (v/v) FBS, 50% (v/v) snail serum, or DOPC (125 µg/mL at 4 hours or 62 µg/mL at 14 hours). The data in panels L–P and the data in panels B–K were collected in separate experiments on separate days. (L) The sphericity of DOPC-induced aggregates was similar to FBS- and snail serum-induced aggregates at 4 hours and lied between those at 14 hours. (M) Density of DOPC-induced aggregates were slightly less than snail serum-induced aggregates at both time points. (N) DOPC-induced aggregates were similar in roughness to snail serum-induced aggregates, both of which were rougher than FBS-induced aggregates at both time points. (O) There were significantly more non-aggregated cells present in DOPC-induced aggregate images than in either serum-induced sample at both time points. (P) DOPC-induced aggregates were similar in height to FBS-induced aggregates and shorter than those induced by snail serum. (Q panel in top-right) The average aggregate area monitored every 20 minutes for 36 hours after induction by 62.5 µg/mL of DOPC vesicles over time. For plots, means ± sem (n=3) are shown, and values from individual replicates are displayed with small circles.

Finally, we also monitored the aggregation dynamics induced by DOPC over time. We found that the dynamics of DOPC-induced aggregation were similar to the case of snail serum induction, although they formed quicker and dissipated earlier (**Fig. 4Q** [top right panel] and **Movie S2**).

### *Capsaspora* responds to host infection with *Schistosoma mansoni*

Since aggregation was sensitive to the concentration of lipid inducers, we hypothesized that *Capsaspora* could sense physiological changes in its host snail that alter serum lipid levels. If so, its aggregation may be an adaptive response to these changes in its host physiology. Some lipids present in *Biomphalaria* serum are known to change in response to the snail’s metabolic state. For example, snails fed different diets have exhibited different serum levels of triglycerides and other neutral lipids.^38,39^ Furthermore, infection with parasites can alter lipid levels in *Biomphalaria* snails. For instance, serum cholesterol and triglyceride levels decrease in *B. glabrata* in response to infection with *Echinostoma paraensei*.^40^ Triglycerides in the entire snail body also drop in response to infection with *Echinostoma caproni*.^41^ Particularly of interest for our DOPC-induced aggregation, whole-snail PCs have been shown to double after 8 weeks of infection by *E. caproni*.^41^ Therefore, to test if *Capsaspora* could sense schistosome-induced differences in host serum lipid composition, we harvested serum from naïve outbred M-line *B. glabrata* snails as well as identical M-line snails that had been infected with 10 PR1 (Puerto Rico Strain 1) *Schistosoma mansoni* miracidia (and were shedding mature schistosome cercaria). We tested the two sera for *Capsaspora* aggregation induction and observed a substantial difference between them (**Fig. 5A**). Remarkably, serum from infected M-line snails induced *much greater aggregation* than the naïve M-line snails, which showed comparatively little aggregation activity (**Fig. 5A–B**). Moreover, the aggregation induced by serum from infected M-line snails was dose-dependent (**Fig. 5C**).

**Figure 5:**
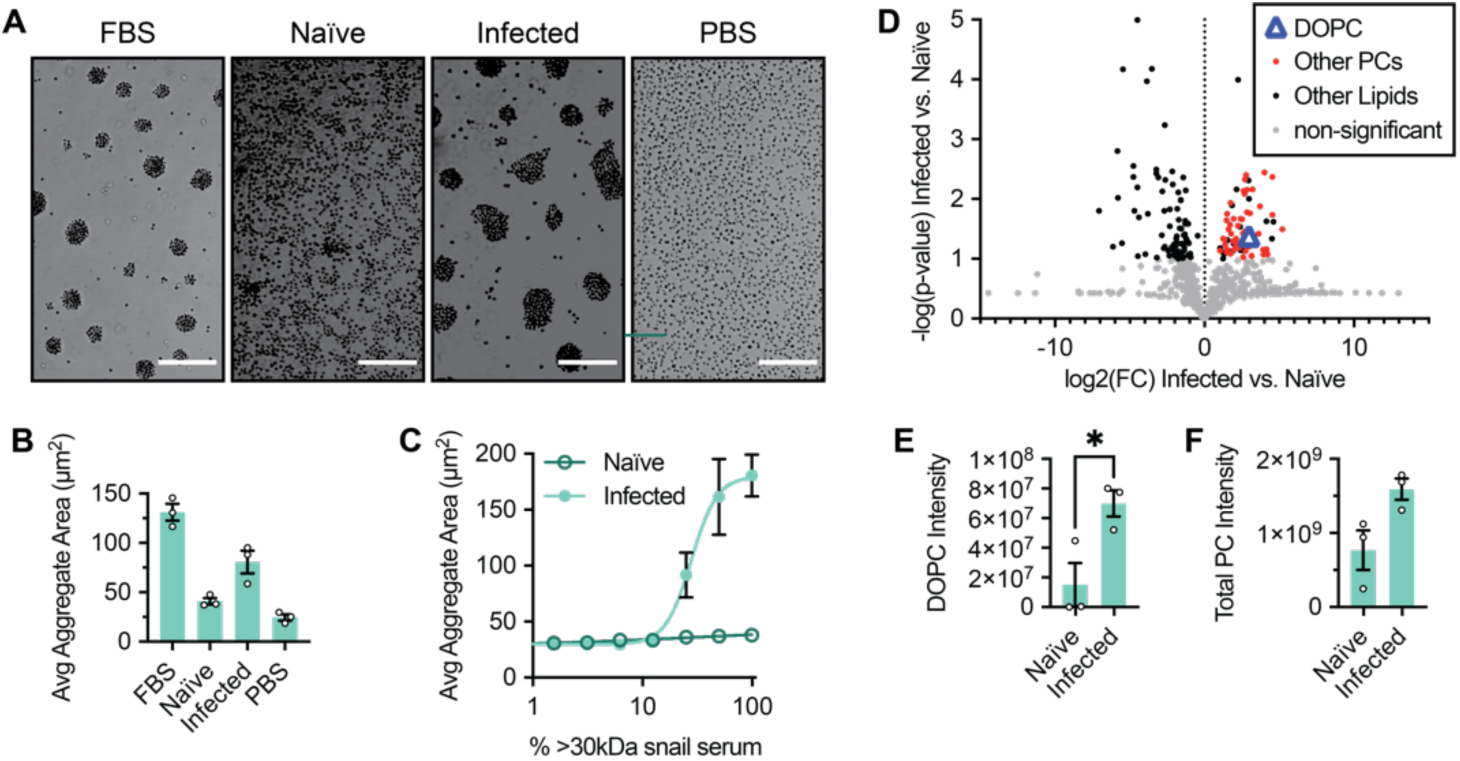
*Capsaspora* aggregates differently in response to serum from snails infected with schistosomes. (A) Representative images of *Capsaspora* cell aggregates induced by either 5% (v/v) FBS, 50% (v/v) >30 kDa serum from naïve M-line snails, 50% (v/v) >30 kDa serum from schistosome-infected M-line snails, or 5% (v/v) PBS control. Scale bars represent 100 µm. (B) Average aggregate area of *Capsaspora* cells induced by snail serum samples compared to FBS and PBS controls. (C) Dose response curve of average aggregate area of cells induced by naïve and infected sera. Infected serum induced aggregation significantly better than naïve snail serum. (D) Volcano plot showing the fold-change (FC) of individual lipids in naïve vs. infected serum, detected by LC-MS/MS. The p-values were calculated from analysis of 3 distinct batches of snails in each condition. Many PCs showed significant increases in the infected snails (see SI for the full table of lipids). (E) Comparison of DOPC [M+H]^+^ intensity in each sample (student’s t-test p=0.03). (F) Comparison of total identified PC [M+H]^+^ intensity summed in each sample (student’s t-test p=0.05). For plots B, C, E, and F, means ± sem (n=3) are shown, and values from individual replicates are displayed with small circles.

To explain the difference, we hypothesized that the infected snail serum contained higher levels of DOPC and/or other phosphatidylcholines (PCs). Thus, we analyzed the lipid contents of each sample using LC-MS/MS. Of the 800 lipids detected, we found 104 to be significantly different between the infected and naïve snail sera (**Fig. 5D, Table S1**). Triglycerides and diglycerides generally decreased upon *S. mansoni* infection, which is consistent with previous studies of echinostome infections.^40,41^ Also consistent with previous work of echinostome-infected snails,^41^ *many of the PC lipids were significantly higher in the infected samples* (**Fig. 5D**), *including DOPC* (**Fig. 5E**). Furthermore, not a single detected PC was significantly depleted in the infected sample. Some abundant PCs were not substantially different between the two samples, rendering the sum of all PCs insignificantly different by a t-test (**Fig. 5F**). However, since several individual PCs were increased in the infected snails, it is plausible that the increased concentrations of a certain class of PCs in the infected snail serum caused the improved aggregation. Overall, these data show that *Capsaspora’s* aggregation phenotype is sensitive to schistosome-induced changes in host serum and that *Capsaspora* aggregation correlates with the serum concentration of DOPC and many other PCs.

### *Capsaspora* responds differently to serum of different *Biomphalaria glabrata* strains

Unexpectedly, we observed that the naïve M-line snail serum above failed to induce robust aggregation at any tested concentration (**Fig. 5C**). This finding contrasted the previous results using serum from naïve NMRI snails, which repeatedly induced robust aggregates (**Fig. 1E–G**). Moreover, this *Capsaspora* aggregation difference between M-line and NMRI snail sera persisted upon repeated inspection (**Fig 6A–B**). This difference could be due to the genotypes of the snails, or it could be due to snail age or different snail husbandry conditions (*e.g.,* temperature, food, or snail density in tanks, see **Materials and Methods** section). To explain the different aggregation potencies of the sera, we hypothesized that they might contain different levels of phosphatidylcholine lipids. Specifically, the NMRI snail serum may have contained higher concentrations of PCs. Thus, we quantified the lipid content of serum from naïve M-line and NMRI snail strains by LC-MS/MS analysis. The two naïve snail sera did not contain significantly different DOPC concentrations (**Fig. 6C–D**). However, several other PCs were significantly higher in the NMRI serum compared to the M-line serum, and no PCs were lower in the NMRI serum (**Fig. 6C**). Although the sum of PCs was insignificantly different between the samples (**Fig. 6E**), it is plausible that a subset of specific PCs is responsible for the observed difference in aggregation induction by our NMRI and M-line snails. Alternatively, we explored other explanations below.

**Figure 6:**
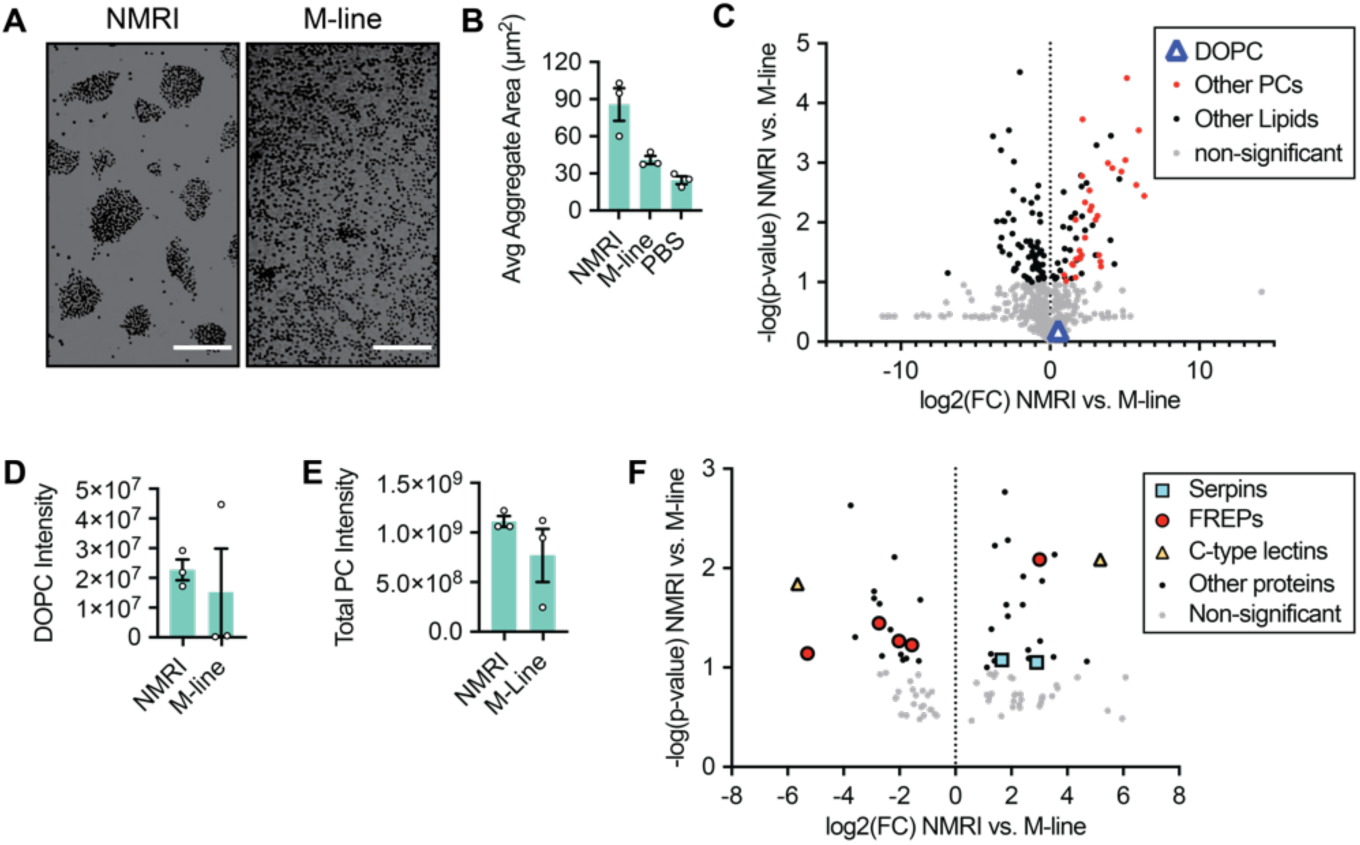
The *Capsaspora* aggregation phenotype discriminates between snail strains. (A) Representative images of *Capsaspora* aggregates induced by 50% (v/v) >30 kDa snail serum from naïve NMRI snails and M-line snails. Scale bars represent 100 µm. (B) Average area of *Capsaspora* aggregates showed that naïve NMRI snail serum is much more active than naïve M-line serum. (C) Volcano plot of the lipids identified by LC-MS/MS analysis of snail serum samples. Many PCs were higher in the active NMRI serum, but DOPC was not significantly different. (D) Comparison of DOPC [M+H]^+^ intensity in each sample (insignificant change by student’s t-test). (E) Comparison of total identified PC [M+H]^+^ intensity summed in each sample (student’s t-test p=0.2). (F) Volcano plot showing fold-change (FC) of individual proteins in naïve NMRI vs. M-line snail sera determined by LC-MS/MS of tryptic peptides. The p-values were calculated from analysis of 3 distinct batches of snails in each condition. 18 proteins were significantly higher in M-line than in NMRI and 23 proteins were significantly lower in M-line than NMRI. Notably, two serpins, five FREPs, and two C-type lectins were significantly different. For plots B, D, and E, means ± sem (n=3) are shown, and values from individual replicates are displayed with small circles.

We hypothesized that different *proteins* in the sera might also account for the different aggregation induction of the two naïve sera. Two scenarios are possible: 1) inhibitory protein(s) that interfere with aggregation are lower in the NMRI serum, or 2) activator protein(s) present in the NMRI snails are needed for robust aggregation. To test these possibilities, we performed proteomics analysis of the two snail strains and found 18 proteins significantly higher in M-line serum and 23 proteins significantly higher in the NMRI serum (**Fig. 6F, Table S2**). Two members of the serpin superfamily of serine protease inhibitors were significantly higher in the NMRI serum, suggesting that proteases might inhibit aggregation and these protease inhibitors curtail that inhibition. Additionally, several fibrinogen-related proteins (FREPs) and a couple C-type lectins were differentially abundant in the two snail sera. A FREP immune protein or lectin could bind to *Capsaspora* (or the lipid inducer itself) and block aggregation. Alternatively, one of these proteins may activate *Capsaspora* by somehow priming the cells to aggregate. Interestingly, subsequent proteomics analysis of the *infected* M-line snail serum also showed one FREP and one C-type lectin at levels similarly low to their levels in NMRI serum, demonstrating multiple inverse correlations between these specific proteins and aggregation induction activity (**Fig. S4, Table S2**).

Further work is needed to confidently conclude if the PC differences are sufficient to explain the aggregation induction differences across host snail sera (or if other serum components inhibit or promote aggregation). Also, further work will determine if the differential aggregation induction is due to genomic differences between NMRI and M-line snails or environmental differences in snail husbandry. Nevertheless, these data clearly show that *Capsaspora* can recognize other host differences beyond infection with schistosomes.

## DISCUSSION

We have discovered that *Capsaspora* (a predator of parasitic schistosomes) can sense and respond to a specific chemical factor within its host snail’s hemolymph. Namely, *Capsaspora* aggregates in response to a snail serum phosphatidylcholine (PC) (**Fig. 7A–B**). This discovery is the first example of a physiological response of *Capsaspora* to its host, which may inform how it colonizes its host and could therein exclude schistosome parasites.

**Figure 7:**
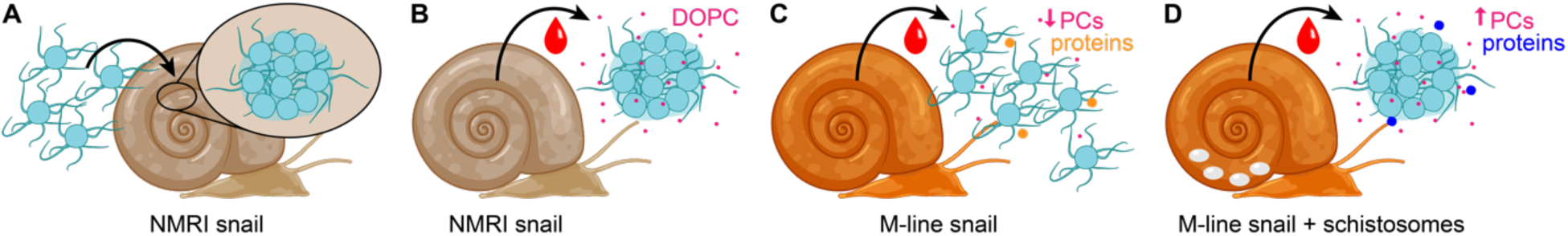
Overview of *Capsaspora* aggregative response to host and upon distinct host-pathogen interactions. (A) When introduced to NMRI snail tissue, *Capsaspora* aggregates. (B) NMRI snail serum also induces *Capsaspora* aggregation, and the serum lipids are responsible for the aggregation—particularly dioleoyl phosphatidylcholine (DOPC). (C–D) M-line snail serum fails to induce robust *Capsaspora* aggregates *in vitro* unless the snails have been pre-infected with schistosomes. Schistosome infection increases the concentration of DOPC (and other PCs) and alters the serum proteome.

*Capsaspora* was recently shown to aggregate in response to FBS *in vitro*.^27^ However, mammalian serum is irrelevant to the expected natural life of *Capsaspora*, which has only been detected in *Biomphalaria* snails and once in fish feces (possibly caused by the fish eating snails).^42^ Thus, our discovery that *Biomphalaria* tissue and hemolymph induce aggregation suggests that this aggregative phenotype *is an ecologically relevant response to a host environment*. As further support for the ecological significance of this phenotype, we found that all three existing isolated strains of *Capsaspora* aggregate in response to snail hemolymph—suggesting that *Capsaspora* aggregation is a widely conserved cellular response to the host environment.

We furthermore identified a single pure lipid from the snail hemolymph that is sufficient to induce aggregation: dioleoyl phosphatidylcholine (DOPC). Since phosphatidylcholines are major components of LDLs, this finding is consistent with our earlier work, which revealed that a combination of LDL lipids collectively induced aggregation.^27^ Quantification of DOPC in snail serum revealed that its concentration is below the threshold level required for pure DOPC vesicles to quickly induce aggregation. Therefore, it is likely that other components in snail serum promote aggregation, as well. For example, other phosphatidylcholines may also induce aggregation—we are currently assessing the aggregation induction ability of a wide panel of phospholipids. Additionally, other serum components may synergize with DOPC to increase its potency. In fact, the aggregate morphologies upon DOPC induction differed some from those induced with whole snail serum (or with FBS), which also suggests that additional serum components contribute to aggregate formation. Since some *Capsaspora* mutants have exhibited similar differences in aggregation morphology, the unknown serum components may interact with these newly characterized pathways in *Capsaspora* to modify aggregate structure.^37^

Having identified the molecular cue of aggregation induction in snail serum, we asked whether *Capsaspora* would be able to use this cue to sense changes in the physiological state of its snail host. Other work has shown that certain lipid classes exhibit different abundance in snails under starvation and infection.^38–41^ Remarkably, we found that schistosome-infected snails harbored elevated concentrations of DOPC (the aggregation inducer), and correspondingly, this serum more potently induced *Capsaspora* aggregation (**Fig 7C–D**). Therefore, it appears that *Capsaspora* can sense the infected state of its host snail and responds with more robust aggregation.

Furthermore, we discovered that *Capsaspora* can differentiate between different inbred strains of snails grown under different laboratory conditions. It aggregated far better in uninfected NMRI snail serum than in uninfected M-line serum. Surprisingly, we found no significant difference in the concentrations of DOPC between the two snail strains. However, other PCs were significantly higher in NMRI serum (and none were higher in M-line serum). Therefore, the induction ability may still rely on a threshold concentration of certain PCs. Alternatively, aggregation may be induced or inhibited by other factors that differ between the strains. The serum proteomes of the two snail strains revealed different levels of FREPs. These proteins serve as immune effectors in snails^43^ and are known to differ in expression across inbred lines^44^ and during infections.^45^ M-line snails may produce specific FREPs that block *Capsaspora* aggregation by preventing its interaction with DOPC or by directly inhibiting its cell-cell contacts. If this hypothesis proves true, it is notable that some FREPs were also depleted in the infected M-line serum, including one that was depleted in the active NMRI serum. Therefore, the increased aggregation in infected serum could be *both* due to increased DOPC concentrations *and* decreased concentrations of certain FREPs. Alternatively, the NMRI FREPs may play a role in promoting DOPC-induced aggregation.^46^ Serpin protease inhibitors, a class of proteins that have previously been investigated for their role in host-pathogen interactions,^47^ were also increased in the aggregation-inducing NMRI serum, possibly indicating an anti-aggregation effect by proteases. Ultimately, the chemical composition of sera from different snail strains grown in different laboratory conditions yielded remarkably different *Capsaspora* aggregation, indicating a means of distinguishing between potential hosts.

In total, these discoveries raise the question of why *Capsaspora* aggregates inside its snail host. Furthermore, it is curious what benefit *Capsaspora* would gain from aggregating more in schistosome-infected snails than in naïve snails. The potential benefits of multicellular forms are myriad,^48–50^ but we mention a few specifically here. First, diverse microbes form multicellular adhesive phenotypes to remain in a favorable environment.^48–50^ For *Capsaspora*, the large aggregates may localize to certain favorable tissues or avoid excretion from the host. A second potential benefit is efficient utilization of secreted exoenzymes. Like many osmotrophic microbes, *Capsaspora* is believed to secrete enzymes to liberate soluble nutrients.^28^ As more cells co-localize, they benefit from each other’s “common goods” and feed more efficiently. This logic has explained the frequent exoenzyme regulation by quorum sensing (or “diffusion sensing”) in bacteria.^51^ Finally, aggregation may protect *Capsaspora* from the snail immune system. Multicellular growth is known to protect symbiotic and free-living microbes from predation.^52–55^ Because snail hemolymph contains immune cells (hemocytes) that can engulf prey^56^ as well as release toxic soluble factors,^57,58^ *Capsaspora* aggregation may afford protection from these insults. Why *Capsaspora* would particularly benefit from aggregation in schistosome-infected snails is also unclear. Perhaps the elevated PC levels indicate a more nutrient-rich environment that favors adhesion and cooperative feeding. Alternatively, higher PC concentrations may indicate an elevated immune state that necessitates protective aggregation.

It is tempting to speculate that the different aggregation responses to sera from different snails will lead to different abilities of *Capsaspora* to colonize those snails. Because serum-induced aggregation is conserved across all existing isolates of *Capsaspora*, the phenotype is likely important for its natural fitness in the snail. However, further work is necessary to determine the necessity of aggregation for snail colonization. If aggregation indeed promotes colonization, then *Capsaspora* may fare better in schistosome-infected snails and in NMRI snails than in uninfected M-line snails. In ongoing work, we are testing this hypothesized dependence of *Capsaspora* colonization on snail strain and infected status.

Lastly, aside from its symbiosis with snails, *Capsaspora* is also an important model for studying the evolution of multicellular phenotypes in animals.^23^ It is one of the closest living relatives of animals and contains many genes that are important for cell-cell adhesion and signaling in animals.^59^ Furthermore, Ruiz-Trillo and co-workers have proposed that *Capsaspora*-like aggregation in the unicellular ancestor of animals may have been a pivotal step in the evolution of the first multicellular animals.^60^ Specifically, cell aggregation may have been the first step to establish obligate multicellular animals (instead of, or in addition to, incomplete separation of cells after clonal cell division). Therefore, dissecting how and why aggregation occurs in *Capsaspora* may illuminate potential mechanisms of aggregative multicellularity in the unicellular ancestor of animals and in the earliest animals. Now that a pure chemical inducer is known, it can be leveraged into biochemical and genetic experiments to determine how *Capsaspora* regulates its multicellularity, both *in vivo* and *in vitro*. Comparative genomics of the regulatory pathway(s) across animals and non-animal holozoans can then assess the potential ancestry of regulated cell-cell adhesion phenotypes in animals. Along this line, it will also be informative to determine if *Capsaspora’*s relatives (*e.g., Pigoraptor* spp.^61^, *Ministeria vibrans*^62^*, Txikispora philomaios*^63^, and *Tunicaraptor unikontum*^64^, see **Fig. S5**) also aggregate in response to PCs, especially given that they exhibit free-living or parasitic distinct lifestyles. If so, this response to PCs may be ancestral, predating *Capsaspora*’s symbiosis with snails and perhaps even predating the divergence of animals from their unicellular holozoan ancestor. Overall, by more deeply elucidating the mechanism of this lipid-induced aggregative response and the breadth of its conservation across phyla, we (and others) can discern the significance of lipids in regulating multicellular behaviors in the evolution of animals.

## CONCLUSION

In sum, we discovered that a snail symbiont that is capable of killing human parasites can adapt its behavior to its host snail environment. Namely, the symbiont *Capsaspora* forms multicellular aggregates in *B. glabrata* snail host tissue and hemolymph. *Capsaspora* senses at least one specific lipid (DOPC) from its host serum and likely uses the concentration of this lipid and its analogs to sense the physiological state and genotype of its host. Because this response is conserved across several symbiont isolates, it is likely significant for the natural life of *Capsaspora*. These findings pave the way for further work to investigate the persistence of *Capsaspora* within the host, the importance of *Capsaspora* aggregation for its ability to colonize snails, and ultimately, this symbiont’s ability to limit the proliferation and spread of parasitic schistosomes from snails.

## ACKNOWLEDGMENTS

We thank the Schistosomiasis Resource Center for provision of snails and schistosomes. NMRI snails were provided by the Schistosomiasis Resource Center of the Biomedical Research Institute (Rockville, MD) through NIH-NIAID Contract HHSN272201700014I. NIH: B. glabrata (NMRI). We thank Margaret Mentink-Kane and André Miller for instruction on rearing snails and collecting hemolymph. We thank Timothy Yoshino for advice and initial provision of snail serum. We thank Eric Loker and Chris Bayne for advice. We thank the Light Microscopy Center at Indiana University for support in image acquisition and analysis (funding provided by the NIH grant NIH1S10OD024988-01). We also thank the Indiana University Nanoscale Characterization Facility, Electron Microscopy Center, and Laboratory for Biological Mass Spectrometry for use of their instruments. We thank Jon Trinidad for proteomics assistance and John Asara for lipidomics assistance. We thank Pranav Danthi for use of the Incucyte imager and Andrew Zelhof for assistance in injecting *Capsaspora* into snails. We thank Jonathan Phillips for advice with *Capsaspora* transfection. We thank Iñaki Ruiz-Trillo for feedback on the manuscript. We also thank the entire Gerdt lab for insights and support that helped advance this project. This work was supported by the National Institutes of Health (R35GM138376) to J.P.G., as well as NIH grants (R37AI101438, P30GM110907). R.Q.K. was supported by an NIH training grant (T32GM131994). The content of this paper is solely the responsibility of the authors and does not necessarily represent the official views of the National Institutes of Health.

## AUTHOR CONTRIBUTIONS

Conceptualization, R.Q.K., N.R.R., J.P.G.; Methodology, R.Q.K., M.R.L, C.G., J.P.G.; Investigation, R.Q.K., E.B.G., M.R.L., M.C.S., C.G., L.P.B.; Writing – Original Draft, R.Q.K., J.P.G.; Writing – Review & Editing, R.Q.K., M.R.L., N.R.R., J.P.G.; Visualization – R.Q.K., J.P.G.; Supervision, J.P.G.; Funding Acquisition, J.P.G.

## MATERIALS AND METHODS

### Cell strain and growth conditions

*Capsaspora owczarzaki* cell cultures (strain ATCC®30864) were grown axenically in 25 cm^2^ culture flasks with 6 mL ATCC media 1034 (modified PYNFH medium) containing 10% (v/v) heat-inactivated Fetal Bovine Serum (FBS), hereafter *growth media*, in a 23°C incubator. Adherent stage cells (filopodiated amoebae) at the exponential growth phase were obtained by passaging ∼100–150 µL of adherent cells at ∼90% confluence in 6 mL of growth media and grown for 24– 48 hours at 23°C until ∼100% confluent.

M-line *Capsaspora owczarzaki* cell cultures (strain ATCC®50973) were grown axenically in 25 cm^2^ culture flasks with 6 mL *growth media*, in a 23°C incubator. Cells were maintained by passaging ∼1 mL of adherent cells at ∼1x10^6^ cells/mL in 5 mL of growth media and grown for 1 week at 23°C until ∼100% confluent.

Salvador *Capsaspora owczarzaki* cell cultures (strain ATCC®50974) were grown axenically in 25 cm^2^ culture flasks with 6 mL *growth media*, in a 23°C incubator. Cells were maintained by passaging ∼3 mL of adherent cells at ∼2.8x10^6^ cells/mL in 3 mL of growth media and grown for 1 week at 23°C until ∼100% confluent.

#### Snail rearing and maintenance conditions

*Biomphalaria glabrata* NMRI snails were obtained from the Biomedical Research Institute (Rockville, MD) (BRI) Schistosomiasis Resource Center. Snails were kept in tanks with about 5 L of artificial pond water (BRI protocol: 0.46 µM FeCl3, 220 µM CaCl2, 100 µM MgSO4, 310 µM KH2PO4, 14 µM (NH4)2SO4 in water adjusted to pH 7.2 with NaOH) with no more than 50 snails per tank until experiments were conducted. Snails were fed romaine lettuce once per week or earlier if they ran out unless otherwise stated. Pond water was changed once per week, or sooner if water was cloudy, by transferring all snails into a new tank with fresh artificial pond water.

*Biomphalaria glabrata* outbred M-line snails were maintained in plastic 20 L tanks filled with 15 L of artificial pond water with no more than 30 snails per tank. Snails were fed red leaf lettuce and 2 Wardly® shrimp pellets 2 times a week. The water was changed once per month. Snails were maintained between 25-27°C on a 12h:12h light-dark cycle.

#### Harvesting snail serum

Serum was harvested from snails measuring between 10 and 30 mm. Snails were removed from their growth tanks, rinsed with autoclaved water, and dried with paper towels or KimWipes. Serum was harvested by the headfoot retraction method.^65^ Briefly, using 200 µL micropipettes or 1 mL glass pipettes, the tip was tapped gently onto the snail headfoot, causing it to retract and hemolymph to pour out from the hemal pore. As the foot retracted, serum was collected into the pipette tip and transferred to microcentrifuge tubes. After harvesting all possible serum, snails were placed in stage 1 of a 2-stage euthanasia solution^66^ (95% (v/v) water + 5% (v/v) ethanol) for ten minutes, and then transferred to stage 2 (95% (v/v) ethanol + 5% (v/v) water) for five minutes. Tubes containing snail serum were centrifuged at 14,000xg for 15 minutes to move any mucus collected to the top of the tube. Bright red, transparent serum was collected from the bottom of the tube and diluted 1:1 in Chernin’s balanced salt solution (CBSS+)^67^ before sterile filtering through at 0.22 µm filter. Sterile serum was stored at 4°C until use.

### General aggregation assay methods

All aggregation assays were performed at room temperature. Brightfield imaging was performed using the following instruments: Leica DMi1 inverted microscope with an MC120 HD camera, Leica DMIL inverted microscope with Flexacam C3 camera, an Olympus OSR spinning disk confocal microscope with a Hammamatsu Flash 4 V2 camera, and an Incucyte S3 Live-Cell Analysis System. Depending on well size and microscope used, each well was imaged at up to 3 distinct locations using 5X or 10X magnification. Average aggregate areas were typically measured by batch processing with a standard macro script in Fiji Imaging Software^27,68^ (see *Image analysis* below).

#### Aggregation assay on ultra-low attachment plates

Two days before the assay, 100% confluent adherent cells growing in 25 cm^2^ culture flasks were given fresh growth media (termed the “feed step”). One day before the assay, cells were washed and resuspended in FBS-free assay media and allowed to sit overnight (termed the “starve step”). After starvation, the day of the assay, 8*10^5^ cells were seeded in 180 µL of FBS-free media per well in a 96-well ultra-low attachment microplate (#CLS3474, Corning) and allowed to settle for 2 hours. Putative aggregation inducers were added such that the total volume in a well was 200 µL. Typically, aggregates were assessed by microscopy after 90 min.

#### Image analysis for aggregation assays

Average aggregate areas were typically measured by batch processing with a standard macro script in Fiji Imaging Software version 2.1.0/1.53c.^27,68^ Briefly, the macro steps included: set the scale of the image appropriate for the microscope conditions, convert the image to binary, analyze particles (size 0-infinity), export results to clipboard. A copy of the FIJI macro is available upon request.

### *Capsaspora* forms multicellular aggregates in snail tissue

#### Generating a *Capsaspora* line stably expressing tdTomato (related to Fig. 1C–D, I–M, and Fig. 4A–P)

A tdTomato expression plasmid was generated from plasmid pJP72.^26^ The tdTomato gene was synthesized by GenScript with codons optimized for *Capsaspora*. The gene was cloned into pJP72, replacing the mScarlet protein-coding region. The resulting plasmid (pJG01) is available from Addgene (#213505), and its sequence is deposited there (www.addgene.org). *Capsaspora* cells were transfected following the protocol by Phillips et. al (2022).^26^ Briefly, on the first day, 3*10^5^ cells in exponential growth phase were seeded in 800 µL onto sterile 12 mm circular glass coverslips in a well of a 24-well plate and allowed to settle overnight. On the second day, the growth medium was removed and replaced with transfection medium (Scheider’s Drosophila Medium with 10% (v/v) FBS, supplemented with 25 µg/mL ampicillin) and allowed to incubate for 10 minutes. Two samples each were treated with Opti-MEM with 2 µg of transfection DNA (either pJG01, or negative control without DNA) along with TransIT-X2 transfection reagent (Mirus Bio) premixed and allowed to incubate 5 minutes at room temperature. Cells were treated with 70 µL of transfection mix and incubated at 23°C for 24 hours. On the third day, the transfection medium was removed and replaced with standard growth medium, and cells were allowed to recover for 24 hours. On the fourth day, growth medium was removed and replaced with medium supplemented with selective drug (Geneticin at 320 µg/mL). Transfected cells were grown in selective medium for two weeks changing media with fresh selective media every 3 days. Cells in the negative control wells were dead after one week of drug selection. Red fluorescence from tdTomato-positive cells was screened on an Olympus spinning disk confocal microscope using a 561 nm laser, which confirmed >99% of cells expressed tdTomato. After two weeks, the culture of cells was expanded from a 24-well plate to 25 cm^2^ culture flasks and maintained in selective growth media until further use.

#### Injection of fluorescent *Capsaspora* into snail tissue (related to Fig. 1C–D and Fig. S1C–D)

Snails were prepared and mounted on microscope slides according to the BRI protocol.^69^ Briefly, snails measuring between 10 and 20 mm were placed in warm artificial pond water (70°C) for 50 seconds, then immediately submerged into a cold water bath for 60 seconds. Snails were removed from their shell by gently pulling on the foot with forceps and placed on microscope slides for injection. Snails were injected using glass capillary needles (World Precision Instruments, 1B100F-6) pulled to an opening size of approximately 100 µm with a Model P-87 Sutter Instrument Micropipette Puller. Two days before the experiment, a 100% confluent flask of *Capsaspora* cells stably expressing tdTomato were “fed” by replacing the selective growth media with fresh selective growth media. The day before the experiment, the *Capsaspora* cells were washed and resuspended in FBS-free media containing no antibiotics and allowed to starve overnight. On the day of the experiment, cells were washed and then resuspended with Chernin’s balanced salt solution (CBSS+)^67^ before injection. Snails were injected with 20 µL of *Capsaspora* cells suspended in balanced salt solution (4*10^7^ cells/mL) into their mantle cavity or pericardium. After injection, needles were left in the tissue for several minutes to allow the hemolymph to clot and prevent *Capsaspora* from bleeding back out. To determine if aggregation was calcium dependent, the calcium chelator EGTA was pre-mixed with cells to a final concentration of 250 mM before injecting 20 µL into snails. As a negative control, cells were injected through the needle directly onto a microscope slide with no snail. Also, snails that were never injected were imaged (no red fluorescent cells of the correct size were observed). Red fluorescence was imaged using an Olympus spinning disk confocal microscope with a 547 nm excitation laser. Brightfield and green fluorescence (488 nm) images were taken as well to see surrounding snail tissue structure.

### *Capsaspora* forms multicellular aggregates in response to macromolecules in host snail serum

#### *In vitro* analysis of snail serum aggregation (related to Fig. 1E)

The standard aggregation assay on ultra-low attachment plates was used to assess activity of 100% (v/v) snail serum compared to 5% (v/v) FBS or 5% (v/v) 1X PBS. To test snail serum, FBS-free assay media was removed by aspiration and replaced with 100% 1X snail serum harvested by the headfoot retraction method. As controls, FBS-free assay media was removed and replaced with either media containing 5% (v/v) FBS or 5% (v/v) 1X PBS. Images of triplicate assay wells were taken every 30 minutes and example images are shown from T-90 minutes.

#### Snail serum fractionation (related to Fig. 1F–G)

Snail serum was fractionated using Amicon Ultra 30 kDa cutoff filters (Sigma, # UFC5030) according to the manufacturer’s directions. Briefly, the filter was first washed with 500 µL of 1X PBS by centrifugation at 14,000xg for 15 minutes. Then 500 µL of harvested serum was added to the cutoff filter and centrifuged at 14,000xg for 15 minutes. The <30 kDa fraction was collected while the >30 kDa fraction was washed three more times with 1X PBS. The standard aggregation assay on ultra-low attachment plates was used to assess activity of 50% (v/v) whole unfractionated snail serum, 50% (v/v) >30 kDa snail serum, and 50% (v/v) <30 kDa snail serum, compared to 5% (v/v) FBS or 5% (v/v) 1X PBS. A dilution series of >30 kDa snail serum was also tested using this method. Images of triplicate assay wells were taken every 30 minutes and analysis was performed on images from T-90 minutes. Average aggregate areas were measured by batch processing with the FIJI macro script reported above.

#### Multiple strains of *Capsaspora* aggregate (related to Fig. 1H)

A non-standard aggregation assay using orbital agitation was set up. *Capsaspora* cells (ATCC 4*10^6^ cells/mL, M-line 8*10^6^ cells/mL, Salvador 8*10^6^ cells/mL) were washed and resuspended in growth media *without FBS,* hereafter referred to as *assay media,* and seeded into a 24-well plate (ATCC 2*10^6^ cells per well, M-line 4*10^6^ cells per well, Salvador 4*10^6^ cells per well). Aggregation inducers were added: 10% (v/v) FBS, 50% (v/v) >30 kDa snail serum, or 10% (v/v) PBS as a negative control), and the plate was agitated at 50 rpm overnight (Celltron Bench-Top Shaker, INFORS-HT). Aggregates were assessed by microscopy after ∼11.5h. Average aggregate area was measured with the FIJI macro script described above.

#### Morphological analysis of tdTomato expressing *Capsaspora* cells after 4 hours of induction (related to Fig. 1I–O, Fig. 4A–P, and Fig. S2A/B/E)

The standard aggregation assay was run using tdTomato expressing *Capsaspora* cells. Aggregates were induced with either 5% (v/v) FBS or 50% (v/v) >30 kDa snail serum, or a dilution series of pure DOPC vesicles (preparation described below) and 3D z-stacks were taken of each replicate well monitored every 4 hours for 48 hours using a Cytation C10 imager (Agilent) with confocal 546 nm excitation laser, laser autofocus, and a 20x objective. Three-dimensional z-stacks from the time point after 4 hours of induction were analyzed using the Imaris batch processing macro described below.

#### Morphological Analysis of Propidium Iodide-stained Aggregates 14 hours after induction (related to Fig. 1I–O, Fig. 4A–P, and Fig. S2C/D/F)

*Capsaspora* aggregates were prepared following the standard aggregation assay protocol in ultra-low attachment plates. Aggregates were induced with either 5% (v/v) FBS or 50% (v/v) >30 kDa snail serum, or a dilution series of pure DOPC vesicles (preparation described below). About 14 hours after induction of aggregates, cells in assay plates were fixed with 4% (v/v) formaldehyde for 30 minutes. Fixed aggregates were washed three times with 1X PBS and then stained with 0.02 mg/mL propidium iodide (PI) for 1 hour. Stained aggregates were washed three times with 1X PBS to remove excess PI before imaging. Cells were imaged using an Olympus spinning disk confocal microscope with 546 nm excitation laser. Three-dimensional z-stacks of aggregates were analyzed using the Imaris batch processing macro described below.

#### Image analysis of 3D confocal experiments (related to Fig. 1I–O, Fig. 4B–P, and Fig. S2)

Three-dimensional z-stacks of aggregates were analyzed by batch processing with Bitplane Imaris Imaging Software version 10.0.1. Briefly, the batch protocol was as follows: create Surfaces, surface grain size = 3 µm, manual threshold 82.114, add Spots, estimate diameter 3.51 µm, background subtraction, “Quality” above automatic threshold, classify spots based on distance to surfaces, threshold 2.91 µm (inside surface), export statistics: surface sphericity, surface intensity inside, number and classification of spots, surface area, surface volume, surface bounding box oriented dimensions. A copy of the Imaris batch protocol is available upon request.

#### Aggregation dynamics over time (related to Fig. 2A–C, Fig. 4Q, and Movie S1–2)

The standard aggregation assay on ultra-low attachment plates was used to assess activity of 5% (v/v) FBS, 50% (v/v) >30 kDa snail serum, or 63 µg/mL DOPC vesicles over time. The corresponding volume of 1X PBS was used as a negative control. Plates were imaged every 20 minutes using the 4X objective on an Incucyte S3 Live-Cell Analysis System. Aggregates were induced 1 hour after initiating Incucyte reads, and plates were kept at 23°C by the system while incubating between images. Resulting image stacks from triplicate wells were first converted into binary and average aggregate area, as well as average aggregate circularity (calculated as the ratio of *x* and *y* bounding box dimensions), was calculated by batch processing with a macro script in FIJI adapted to analyze time stacks. Briefly, the macro is set to: set scale, make binary, gaussian blur (sigma = 4), run “analyze particles” size 3–100000, count, and summarize results. The summary results that were plotted were: average particle area, and circularity (calculated by the ratio of 2D bounding box dimensions). A copy of the FIJI stacks macro is available upon request. Representative binary images from selected time-points are shown.

### Lipids extracted from snail serum are sufficient to trigger aggregation

#### Extraction of lipids from snail serum using butanol and diisopropyl ether (DIPE) (related to Fig. 3A–D)

Snail serum was harvested by the headfoot retraction method.^65^ Lipids were extracted from serum following a protocol from Cham and Knowles (1976).^70^ Briefly, 0.1 mg of EDTA was added to 1 mL of serum in a 4-dram vial. Then, 2.5 mL of premixed 25:75 (v/v) butanol to DIPE was added. The solution was rocked gently at a speed of 50 on a Fisherbrand Digital Platform Rocker for 3 hours. After rocking, the vials were allowed to sit for 15 minutes until the layers separated. The organic layer was carefully removed from the aqueous layer. Note that no proteins in the aqueous layer precipitated. The organic layer was completely evaporated using a Savant SpeedVac SPD2030 at 45°C under 5.1 millitorr of vacuum pressure for 2 hours. Lipids were extracted separately from the sera of three batches of snails and stored at –20°C until use.

#### Preparing lipid vesicles from lipid extracts or from pure lipid samples (related to Fig. 3 and Fig. 4)

Lipids extracted from snail serum samples or commercially obtained (pure synthetic lipids) were redissolved in chloroform in glass vials and transferred to a 1.7 mL microcentrifuge tube. The chloroform was evaporated using a gentle stream of nitrogen and the resulting lipid film was then resuspended in 1 mL each of 1X PBS. The tubes were vortexed for 30 seconds before sonication. Lipid solutions were sonicated on ice with a 50% duty cycle at medium setting for 10 minutes using a single probe attached to a Branson Sonifier Cell-Disruptor 185. During sonication, tubes were kept on ice and tube bottoms were set about 3 mm from the sonicator tip. Lipid solutions went from slightly cloudy to clear after sonication. Sonicated lipids were stored at 4°C for no more than 2 days before use. In the case of crude snail serum lipids (**Fig. 3A-D**), all the lipids were extracted from 1 mL of snail serum, resulting in ∼ 0.3 mg of dried crude lipids. After extracting and drying, crude lipids were reconstituted in 1 mL of PBS buffer and sonicated. Therefore, the extracted lipids were present at their natural serum concentration. The resuspended lipids exhibited similar potency (v/v) as serum.

#### Testing lipid vesicles for aggregation (related to Fig. 3A,E,I,J)

Sonicated lipids were concentrated 10X using an Amicon Ultra 30 kDa cutoff filter and diluted with 1X PBS to the desired concentration before testing. The standard aggregation assay on ultra-low attachment plates was used to test for aggregation induction. Aggregates were induced using the desired lipid concentration (calculated in µg/mL of lipid) and 5% (v/v) of 1X PBS was used as a negative control. Aggregates in triplicate wells were imaged every 30 minutes and analysis was performed on images from T-90 minutes. Average aggregate areas were measured using a macro in FIJI as described above.

#### Transmission Electron Microscope (TEM) imaging of prepared lipid vesicles (related to Fig. 3B,F,K)

Before preparation of samples, Formvar/Carbon film 300 mesh Nickel Grids (Electron Microscopy Sciences FCF300-Ni-25) were ionized using a Pelco easiGlow Discharge System at 0.3 mbar and 15 mAmps for 2 minutes. 10 µL of prepared lipid samples were added to the ionized grids and allowed to sit for 5 minutes. Excess sample was removed from the grids by gently touching Whatman filter paper (Sigma, WHA1001329) to the edge. Immediately after removing samples (before grid dries) 10 µL of 2% (v/v) Uranyl Acetate pre-diluted in water was added to grids and allowed to sit for 10 seconds. Then the stain was removed with the edge of Whatman filter paper. Grids were allowed to dry completely before imaging (about 10 minutes). Prepared grids were imaged using a JEOL-JEM 1010 transmission electron microscope (TEM) with an 80 kV operating voltage, equipped with a 1k x 1k Gatan CCD camera (MegaScan model 794) using a tungsten filament as its electron source.

#### Determination of prepared lipid vesicle size using Dynamic Light Scattering (DLS) (related to Fig. 3 C,G,L)

400 µL of prepared lipid vesicles suspended in 1X PBS at room temperature were transferred to a BRAND UV micro disposable cuvette (BR759200-100EA) and placed in a Malvern Panalytical Zetasizer Nano ZS DLS. Size measurements were recorded in a range of 0.3 nm to 3 mm as raw intensity and then normalized based on particle size using the Malvern software to give the final distribution of particle sizes in each sample. The “number” distribution was plotted and Z-averages (nm) were reported. Triplicate measurements were recorded for each sample.

#### Mass spectrometric analysis of lipid samples (related to Fig. 3D,H, Fig. 5D–F, Fig. 6C–E, Fig. S3, and Table S1)

For most lipidomics analysis, fully dried lipid samples were shipped to the Beth Israel Deaconess Medical Center (BIDMC) Metabolomics Core. The LC-MS/MS based non-polar lipidomics profiling work was conducted by Dr. John Asara at the BIDMC Mass Spectrometry Core in Boston, MA USA. For calculation of the amount of DOPC in snail serum samples, 1 mL of snail serum was spiked with a known volume of SPLASH Lipidomix II mass spectrometry standard (Avanti Polar Lipids, #330709) and then extracted according to the method described above. Dried lipid samples were resuspended in 2:1 (v/v) isopropanol/methanol. High-resolution electrospray ionization (HR-ESI) mass spectra with collision-induced dissociation (CID) MS/MS were obtained using an Agilent LC-q-TOF mass spectrometer 6530 equipped with an Agilent 1290 uHPLC system. Metabolites were separated using a Luna 5 μm C5 100 Å LC column (Phenomenex 00D-4043-E0). Mobile phase A was 95% (v/v) Water, 5% (v/v) Methanol, 0.1% (v/v) Formic Acid, and 5 mM Ammonium Formate. Mobile phase B was 60% (v/v) Isopropanol, 35% (v/v) Methanol, 5% (v/v) Water, 0.1% (v/v) Formic Acid, and 5 mM Ammonium Formate. After initially holding 0% phase B for 5 min at 0.1 mL/min, a linear gradient from 20% phase B to 100% phase B was applied over 40 min with a flow of 0.4 mL/min before holding at 100% phase B for another 5 min with a flow of 0.5 mL/min. Data-dependent acquisition was employed to fragment the top masses in each scan. Collision-induced dissociation was applied using a linear formula that applied a higher voltage for larger molecules (CID voltage = 10 + 0.02 m/z)for metabolite profiling and identification. Mass traces collected in positive mode were analyzed in MassLynx software v4.1 and calculated concentrations were normalized to the known concentration of the internal SPLASH standard.

### *Capsaspora* responds to host infection with *Schistosoma mansoni*

#### Infection of M-line snails with PR1 schistosomes (related to Fig. 5 and Fig. 6)

Two mice were exposed to 150 PR1 (Puerto Rican Strain 1) cercariae from snails originally exposed to BRI-derived miracidia, the resulting miracidia from those two mice were used to expose 120 5-6 mm *B. glabrata* M-line snails to 10 PR1 miracidia. All miracidia displayed positive phototaxis, normal shape, and swimming behavior. Snails were exposed for 2 hours in 12-well cell culture plates and then placed into 4 different 20 L tanks. 60 5-6 mm *B. glabrata* M-line snails were sham-exposed (no parasites) in 12-well cell culture plates for 2 hours and placed into 3 different 20 L tanks as control snails. Experimental snails were fed 3 times a week red leaf lettuce and 2 Wardly® shrimp pellets. Snails were maintained between 25-27°C on a 12h:12h light-dark cycle. After 28, 32 and 38 days post-exposure, snails were checked to determine if cercariae were being released (shedding). Snails were placed in 12-well cell culture plates for 2 hours between 10:00 am to 12:00 pm under light. At 38 days post-exposure, 79 *B. glabrata* M-line snails were shedding cercariae (group 1: n=20, group 2: n=20, group 3: n=20, group 4: n=19). Sham-exposed control snails were in 3 different groups (group 1: n=20, group 2: n=20, group 3: n=11).

#### Measuring aggregation induced by naïve and infected M-line snail serum and naïve NMRI snail serum (related to Fig. 5A–C and Fig. 6A–B)

The standard aggregation assay using ultra-low attachment plates was used to determine aggregation potency of snail serum samples. Serum collected from three separate batches of naïve or infected snails were each concentrated to 5X using an Amicon Ultra 30 kDa cutoff filter. Aggregation was induced by a dilution series of each of the >30 kDa snail serum samples. Assay wells were imaged every 30 minutes and representative images from T90 minutes are shown. Average aggregate area was calculated by batch processing with the standard macro script in FIJI as described above.

### *Capsaspora* responds differently to serum of different *Biomphalaria* strains

#### Proteomics analysis of snail serum samples (related to Fig. 6E, Fig. S4, and Table S2)

Serum harvested from three separate batches of naïve NMRI, naïve M-line, and infected M-line snails via the headfoot retraction method were submitted for untargeted proteomics analysis by the Laboratory for Biological Mass Spectrometry at Indiana University.

