## Supplemental Data 1 for "Host lipids regulate multicellular behavior of a predator of a human pathogen"

Document contains:

Supplementary figures S1–S5

Legends for Movies S1–S2

Legend for Supplementary Table S1–S2

### SUPPLEMENTARY FIGURES

#### Figure S1

A

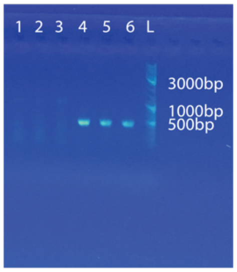


B

**PCR 1 (outer primers)**

| Name | Sequence |
|---|---|
| Capsa_7S | GGACTTTTACTGTGAAAAAA |
| Capsa_28AS | ATCCTTGTTAGTTTCTTTTC |

**PCR 2 (inner primers)**

| Name | Sequence |
|---|---|
| Capsa_5S | GGAGAAAGAAAAAAAAAAGGA |
| Capsa_7AS | CTTGTAGGGGTGAGTTTAGG |

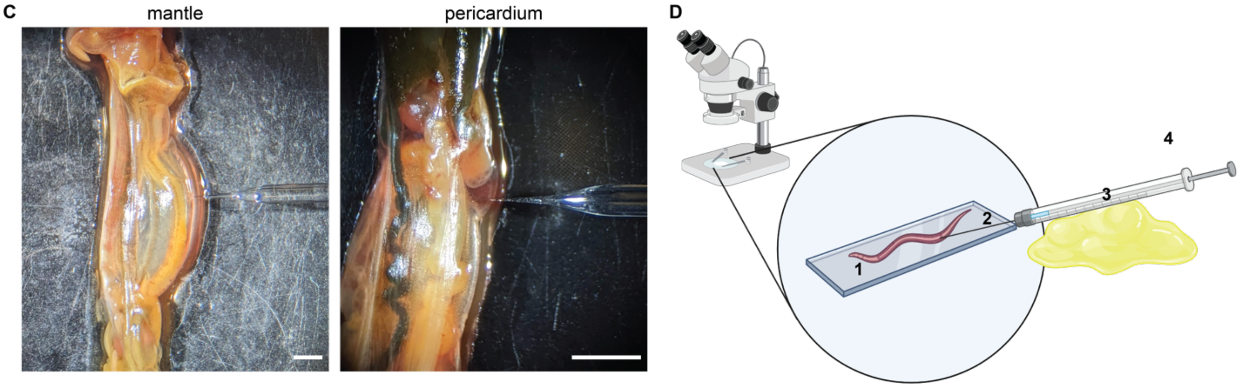


**Fig. S1. *Capsaspora* injections into snails experimental set up.** (A) PCR of *Capsaspora* 18S rDNA in unexposed laboratory NMRI snails (lanes 1–3) and snails that have been exposed to *Capsaspora* for 1 hour (lanes 4–6) showing that *Capsaspora* is not natively present in NMRI snails but is found in exposed snails. (B) Table of primer sequences used for nested PCR protocol for 18S rDNA amplification of *Capsaspora* in snail samples.[1] DNA was extracted from snail tissue using the Macherey-Nagel Nucleospin Tissue kit, and PCR was performed with NEB Q5 polymerase. (C) Representative images of snail injections into the mantle and pericardium. Scale bars are 2 mm. (D) Diagram of snail injection set up: 1. Microscope slide containing a snail pulled from its shell; 2. Glass capillary needle pulled to have a 100 μm opening; 3. Modeling clay for stabilization of injections; 4. 200 μL glass syringe.

**Figure S2**

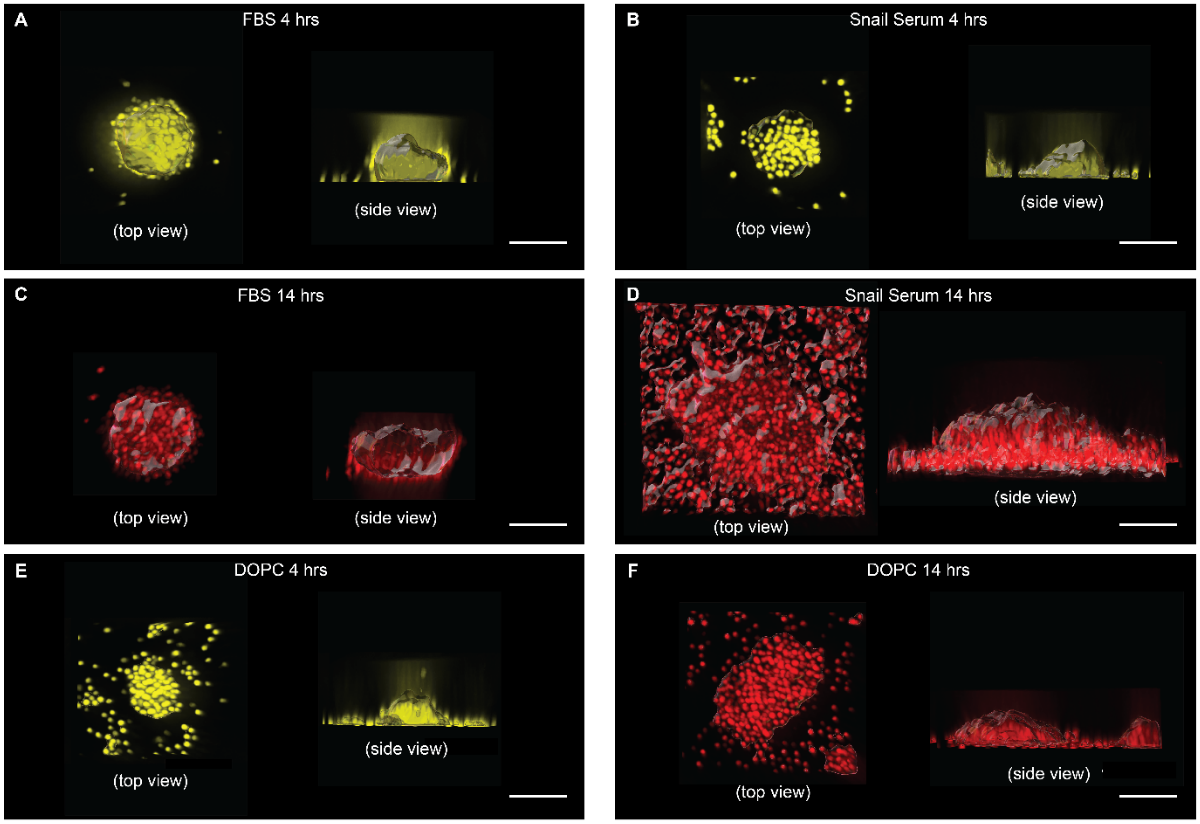


**Fig. S2. Three dimensional images of *Capsaspora* aggregates for morphology analysis.** (A) Top view (left) and side view (right) of representative aggregate induced by 5% (v/v) FBS after 4 hours of induction shows the aggregate is more spherical, dense, smooth, and of medium height. Scale bars are 10 µm. (B) Representative aggregate induced by 50% (v/v) NMRI snail serum after 4 hours of induction shows an aggregate that is very similar to FBS although slightly less spherical and slightly rougher around the edges. Scale bars are 10 µm. (C) Representative aggregate induced by 5% (v/v) FBS after 14 hours of induction shows the aggregate has remained more spherical, dense, smooth, and is taller in height than it was at 4 hours. Scale bars are 10 µm. (D) Representative aggregate induced by 50% (v/v) NMRI snail serum after 14 hours of induction shows aggregates that have remained rough around the edges but have also become less spherical, less dense, and much taller compared to FBS. Scale bars are 20 µm. (E) Representative aggregate induced by 125 µg/mL of DOPC after 4 hours of induction shows the aggregate is of similar shape to FBS and snail serum aggregates but there are many more unincorporated cells. Scale bars are 10 µm (F) Representative aggregate induced by 63 µg/mL of DOPC after 14 hours of induction shows the aggregate is less dense but more spherical than snail serum aggregates, there are also many more unincorporated cells. Scale bars are 50 µm. In all panels, top views are on the left and side views are on the right.

**Movie S1**

*See movie file*

**Movie S1. *Capsaspora* aggregation over time in response to addition of inducers.** (A) Aggregation monitored every 20 minutes for 2 days after addition of 5% (v/v) FBS. (B) or 50% (v/v) >30 kDa NMRI snail serum components. Images were converted to binary in FIJI to enhance contrast. Scale bar is 500 µm.

#### Table S1

*See excel file*

**Table S1. Lipidomics analysis of snail serum samples.** Table shows raw MS/MS intensity data for 800 lipids in NMRI snail serum, naïve M-line snail serum, infected M-line serum, and active vesicles made from the NMRI extracts.

#### Figure S3

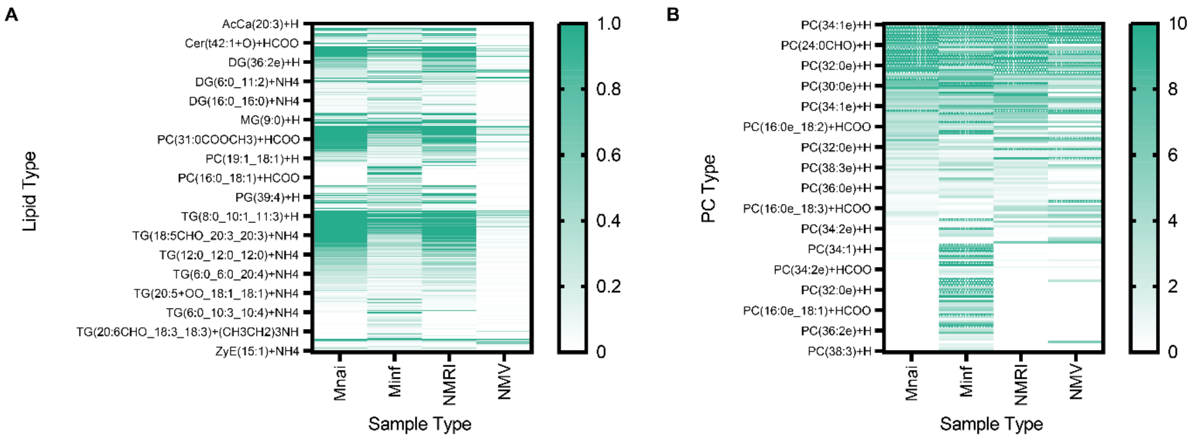


**Fig. S3. Lipidomics analysis of snail serum samples.** (A) Heat maps of all lipid MS/MS intensity data from Table 1 shows general profile of lipids extracted from each sample: naïve M-line snail serum, infected M-line snail serum, naïve NMRI snail serum, and vesicles derived from a 25:75 butanol and di-isopropyl ether extraction of naïve NMRI snail serum. (B) Heat maps of just PC lipids from the same samples as A. For heat maps, gradients of colors represent normalized MS1 intensity detected for each lipid mass (dark = more intense, light = less intense).

#### Movie S2

*See movie file*

**Movie S2. *Capsaspora* aggregation over time in response to addition of DOPC vesicles.** Aggregation monitored every 20 minutes for 2 days after addition of 63 µg/mL of DOPC. Images were converted to binary in FIJI to enhance contrast. Scale bar is 500 µm.

**Table S2**

*See excel file*

**Table S2. Proteomics analysis of snail serum samples.** Table shows raw peptide count data as well as quality scores for proteins identified in NMRI snail serum, naïve M-line snail serum, and infected M-line serum. (Sheet 1 "naive sera") Protein identity, fold change, and p-values for proteins identified in naive NMRI vs naive M-line snail serum samples. (Sheet 2 "inf vs nai M-line") Protein identity, fold change, and p-values for proteins identified in infected vs naive M-line snail serum samples.

**Figure S4**

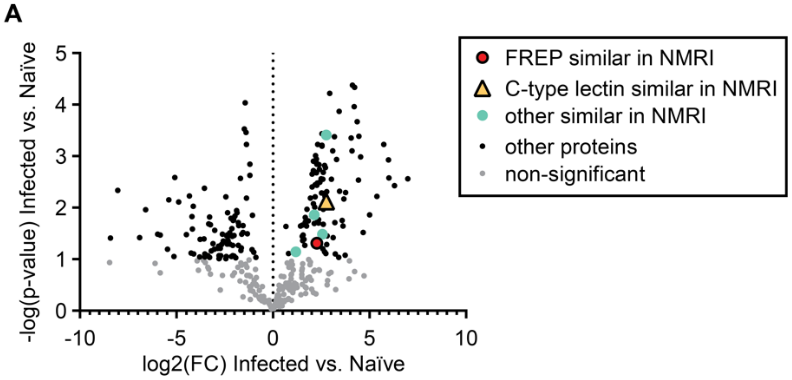


**Fig. S4 Proteomics analysis of infected and naïve M-line snail sera.** (A) Volcano plot of proteins showing fold-change (FC) of individual proteins in infected vs. naïve M-line snail sera determined by LC-MS/MS of tryptic peptides. The p-values were calculated from analysis of 3 distinct batches of snails in each condition. Notably, some differing proteins are inversely correlated with aggregation activity, two proteins were down in both active (infected M-line and naïve NMRI) sera.

**Figure S5**

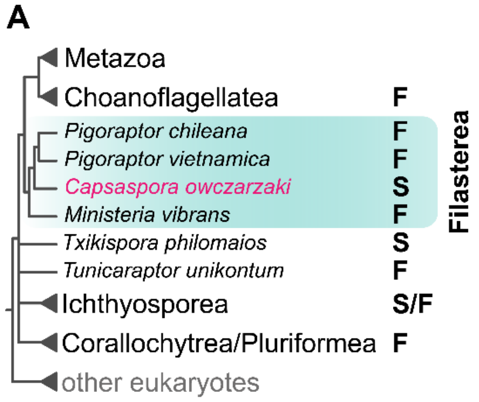


**Fig. S5. Cladogram representing *Capsaspora* and its close unicellular relatives.** Animals (Metazoa) and their closest unicellular relatives, including *Capsaspora* in the Filasterea lineage (turquoise pseudocolor). The phylogenetic relationships of selected taxa are based on several recent phylogenomic studies[2-7]. Uncertain positions are represented with polytomies. Letters depict distinct lifestyles observed in filasterean species and close relatives upon isolation. F: free-living; S: in a symbiotic relationship (*e.g.,* parasitic, commensal, and mutualistic).
